# Phylotranscriptomics provides a treasure trove of flood tolerance mechanisms in the Cardamineae tribe

**DOI:** 10.1101/2024.03.20.585870

**Authors:** Hans van Veen, Jana T. Müller, Malte M. Bartylla, Melis Akman, Rashmi Sasidharan, Angelika Mustroph

## Abstract

Flooding events are highly detrimental to most terrestrial plant species. However, there is an impressive diversity of plant species that thrive in flood-prone regions and represent a treasure trove of unexplored flood-resilience mechanisms. Here we surveyed a panel of four species from the Cardamineae tribe representing a broad tolerance range. This included the flood-tolerant *Cardamine pratensis*, *Rorippa sylvestris* and *Rorippa palustris* and the flood-sensitive species *Cardamine hirsuta*. All four species displayed a quiescent strategy, evidenced by the repression of shoot growth underwater.

Comparative transcriptomics analyses between the four species and the sensitive model species *Arabidopsis thaliana* were facilitated via *de-novo* transcriptome assembly and identification of 16,902 universal orthogroups at a high resolution. Our results suggest that tolerance likely evolved separately in the *Cardamine* and *Rorippa* species. While the *Rorippa* response was marked by a strong downregulation of cell-cycle genes, *Cardamine* minimized overall transcriptional regulation. However, a weak starvation signature was a universal trait of tolerant species, potentially achieved in multiple ways. It could result from a strong decline in cell-cycle activity, but is also intertwined with autophagy, senescence, day-time photosynthesis and night-time fermentation capacity. Our dataset provides a rich source to study adaptational mechanisms of flooding tolerance.

## Introduction

Some plant species can thrive in extreme environments. Underwater conditions are especially challenging to plant life since low gas-diffusion rates hamper oxygen and carbon dioxide (CO_2_) exchange with the environment. This restricts mitochondrial respiration and photosynthesis, respectively, potentially resulting in a severe energy and carbon shortage. Most terrestrial plant species are very flood-sensitive and cannot survive extended periods of flooding. However, the plant kingdom has evolved a variety of adaptive strategies to circumvent these challenges.

One strategy involves plant morphological and anatomical modifications that enhance aeration of submerged plant parts. This includes the development of adventitious roots, aerenchyma formation in stems, petioles and roots, the development of root barriers preventing radial oxygen loss to the anoxic soil (ROL) and shoot elongation to re-establish aerial contact (Pedersen et al. 2021, Combs-Giroir and Gschwend 2024, Lin et al. 2024). In contrast, the quiescence strategy involves restricted growth and metabolism under water until the floods recede, thus conserving valuable resources for regrowth after desubmergence (Voesenek and Bailey-Serres 2013, Combs-Giroir and Gschwend 2024). Among crops, rice cultivars provide the best-studied examples of these contrasting flood-adaptive responses (Xu et al. 2006, Hattori et al. 2009).

Plants that use the above strategies tend to perform best under terrestrial conditions. For heterophyllous species, the favoured environment is less clear. Aerial-formed leaves of these species are like those of terrestrial plants, but those formed underwater are thin (high specific leaf area), have a negligible cuticle, are void of stomates, and exhibit either high dissection or a narrow, elongated morphology. Aerial and aquatic leaves can also vary in CO_2_and bicarbonate usage and carbon fixation pathways (C3, C4, CAM). Heterophylly in amphibious plants optimizes photosynthesis for underwater conditions and prevents desiccation when grown aerially (Van Veen and Sasidharan et al. 2021). These species have been invaluable in unravelling the environmental influence on adaptive leaf development, e.g. *Ranunculus trichophyllus* (Kim et al. 2018), *Potamogeton octandrus* (He et al. 2018), *Callitriche palustris* (Koga et al. 2021), *Hygrophila difformis* (Li et al. 2022), *Rorippa aquatica* (Ikematsu et al. 2023).

Most of these flood-adaptive morphological and anatomical acclimation responses are mediated by the plant hormone ethylene (Sasidharan and Voesenek, 2015, Pedersen et al. 2021, Leeggangers et al. 2023). Being a gas, it quickly accumulates to high concentrations in flooded tissues and is an important flooding stress cue (Voesenek and Sasidharan, 2013). However, ethylene is also a key signal that stimulates senescence, which might not always be beneficial (Rankenberg et al. 2024). Accordingly, species in permanently deluged conditions have lost the ability to produce and/or sense ethylene (Summers et al. 1996, Voesenek et al. 2015, Olsen et al. 2016, Ma et al. 2024).

Most current crops are flood-sensitive with tolerance traits being lost in breeding cycles focused on yield parameters. As climate change exacerbates the frequency and unpredictability of flooding events, it is imperative to intensify our efforts to explore naturally stress-resilient plants. Flood-tolerant plant species serve as valuable models for identifying and understanding tolerance mechanisms, species divergence and adaptation. The Brassicaceae family is particularly interesting in this context, including a range of model, wild and crop species varying in their tolerance to different flooding regimes. Within the Brassicaceae family, a wide range of adaptations to specific water-rich habitats exist (Nikolov and Tsiantis 2017). Most Brassicaceae are sensitive to excess water availability, including the model species *Arabidopsis thaliana* (Mustroph et al. 2009, Lee et al. 2011, Vashisht et al. 2011) or the oil crop *Brassica napus* (Wittig et al. 2021). However, this family also includes species that are flood-adapted. Especially one clade, the tribe *Cardamineae*, contains many flooding-tolerant members, namely from the genera *Cardamine*, *Rorippa*, and *Nasturtium* (Cook 1999).

Our previous work characterized flood responses in *Nasturtium officinale*, watercress, which has adapted to growth in small rivers (Howard and Lyon 1952). This species uses the escape strategy to avoid internal oxygen deficiency conditions (Müller et al. 2021). Within the genus *Cardamine*, the newly emerging model plant species *Cardamine hirsuta* (Kierzkowski et al. 2019, Hajheidari et al. 2019) is not described as flooding-tolerant, but some of its close relatives prefer wet growth conditions (Shimizu-Inatsugi et al. 2017, Akiyama et al. 2021, Kantor et al. 2023). The genus *Rorippa* is also known to prefer wet environments (*R. sylvestris*, *R. amphibia*; Sasidharan et al. 2013, Akman et al. 2014), and one member even shows pronounced heterophylly (*R. aquatica*, Nakamasu et al. 2014, Ikematsu et al. 2023).

In this work, we built on our previous knowledge to further explore flooding resilience mechanisms in Brassicaceae species. We selected two flood-sensitive species, *Arabidopsis thaliana* and *Cardamine hirsute*, and the flood-tolerant *Cardamine pratensis*, *Rorippa sylvestris* and *Rorippa palustris* (Supplemental Figure S1). Despite the variety of adaptive mechanisms, all five species phenotypically exhibited the quiescent strategy. Through a comparative transcriptomic analysis of the shoot response to submergence, we were able to distinguish key players associated with a successful quiescence when flooded. The narrow phylogenetic range, but wide diversity in tolerance found within the Brassicaceae allowed us to quantitively compare species at high genetic resolution. We found distinct routes towards submergence tolerance, which are associated with either a strong downregulation of the cell cycle, or with minimized transcriptomic reconfiguration when under water.

## Material and Methods

### Plant material, growth conditions and stress treatment

Seeds for *Arabidopsis thaliana*, ecotype Col-0, were raised in house. Seeds of *Cardamine hirsuta* were obtained from Miltos Tsiantis. *Rorippa sylvestris* was propagated clonally from plants originally collected in the Netherlands (Stift et al. 2008, Akman et al. 2012). Seeds for the other two species were bought from commercial seed suppliers (*Cardamine pratensis*, www.templiner-kraeutergarten.de; *Rorippa palustris*, www.wildblumen.at).

All plant species were grown on a soil mixture (*A. thaliana* until the 10-leaf-stage, all other species until the 6-leaf stage) about 3-4 weeks after germination, according to established protocols (Müller et al. 2021), under short-day conditions (8 h light, 16 h darkness), 100 µmol photons * m^-2^ * s^-1^ and 23 °C.

Submergence stress was applied to the plants by immersing them in big, transparent boxes (about 50 l volume) filled with 40 l of tap water 24 h before start of the treatment. Treatment was started 2 h after start of illumination. For survival experiments, plants were kept under water for up to 10 weeks. At each time point, 8 plants were removed from the boxes and were kept under normal growth conditions for another two weeks. Surviving plants were counted, while survival was defined as the ability to form new green leaves within the 2-week-recovery period. For molecular and biochemical analysis, the youngest leaves including the meristem were collected after 24 and 48 h of stress treatment, together with aerated controls and frozen immediately in liquid nitrogen.

For measuring growth of the leaf, the length of the youngest leaf was measured with a digital calliper. Measurements were done at the start of the submergence treatment and at three subsequent days. For each leaf, the daily length increment was calculated.

### Biochemical parameters

Frozen tissue (young leaves including meristems) was ground to fine powder, and subsequently used for extraction of metabolites and measurement of soluble sugars by established methods previously described (Riber et al. 2015). Proteins were extracted and activity of ADH was determined as previously described (Gasch et al. 2016).

### RNA extraction, sequencing, and expression analysis

Frozen tissue (young leaves including meristems) was ground and further processed as described in Müller et al. 2021. Subsequent RNA extraction and library preparation was also done exactly as in our previous analysis. Raw reads were processed through adapter trimming of the FASTQ files using the BBDuk algorithm. Detailed approaches of the analysis are described in the Supplemental data. In short, *de novo* transcriptomes were assembled with Trinity version 2.6.6 (Grabherr et al. 2011, Haas et al. 2013) for *C. pratensis*, *R. palustris* and *R. sylvestris*. A genome-guided approach was used for *C. hirsuta* with HISAT2 mapped reads to the genome (Gan et al. 2016, Kim et al. 2019). Sets of orthologous genes between the studied species (Orthogroups) were identified through an all-vs-all discontiguous megablast (Camacho et al. 2009), followed by Orthofinder (Emms and Kelly 2019). Read mapping for expression quantification was done with Kallisto (Bray et al. 2016). Fold changes upon submergence, differences between timepoints and phylogenetic and tolerance-specific effects were determined with the EdgeR package (Robinson et al. 2010). GO enrichment was done with GOseq and the Arabidopsis annotation (Young et al. 2010, Carlson 2019a, Carlson 2019b).

The raw data for this study have been deposited at EMBL-EBI in the European Nucleotide Archive (ENA) under accession number PRJEB73992 (https://www.ebi.ac.uk/ena/browser/view/PRJEB73992) and in ArrayExpress under accession number E-MTAB-13910 (https://www.ebi.ac.uk/biostudies/arrayexpress/studies/E-MTAB-13910).

## Results

### Brassicaceae species vary in their tolerance to complete submergence

To characterize submergence tolerance, the five rosette species were fully submerged at the 10- or 6-leaf developmental stage (vegetative stage ∼3-4 weeks after germination). Plant mortality was periodically checked based on the ability to recover within 10 weeks. *Arabidopsis thaliana* was the most sensitive surviving only 3-4 weeks (Figure 1A, Supplemental Figure S2). The two *Cardamine* species occupy a wetter niche (Table 1, Ellenberg and Leuschner 2010), and accordingly showed better submergence survival. *Cardamine hirsuta* survived about 6 weeks of submergence, while *Cardamine pratensis* survived up to 10 weeks. Both *Rorippa* species survived the 10-week submergence period used here (Figure 1A, Supplemental Figure S2). Extensive algal growth prevented longer submergence periods in our system. Overall, the performance of the five species corresponded to their Ellenberg ecological niche indicator values for moisture levels (Table 1). The survival data indicated a tolerance gradient from a mildly sensitive *A. thaliana* to the moderately tolerant *C. hirsuta*, to the extremely tolerant *C. pratensis*, *R. palustris* and *R. sylvestris*.

**Figure 1.**
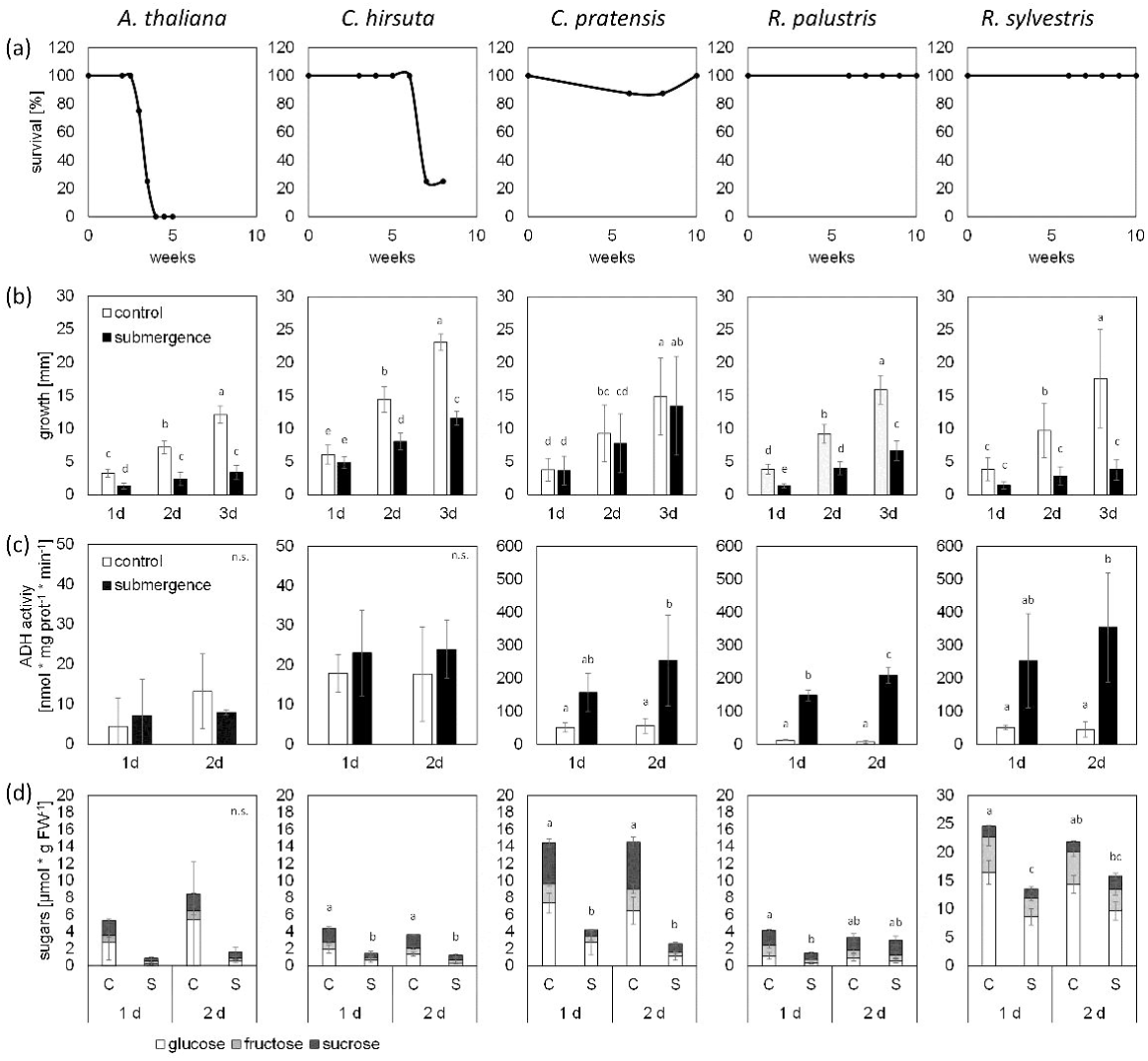
Characterization of submergence tolerance in Brassicaceae. (a) Survival rates under illuminated submerged conditions with regular day-night (8h:16h) rhythm. (b) Growth activity of leaves underwater. Plants with 6 true leaves (*C. hirsuta, C. pratensis, R. palustris, R. sylvestris*) or 10 leaves (*A. thaliana*) were submerged under short-day conditions (black bars) and compared with control plants in air (white bars). The length of the youngest leaf was measured just before the treatment and after 1, 2 and 3 days. The increase in growth is shown with means ± SD from 3 independent experiments with 5 plants per replicate (n = 15). (c) ADH activity under submergence. 3- to 4-week-old plants were submerged under short-day conditions (black bars). After 1 and 2 days of treatment, young leaves were harvested together with leaves from aerated controls (white bars). ADH activities are shown as mean ± SD from 3 to 4 biological replicates. (d) Carbohydrate dynamics in Brassicaceae under submergence. Plants were stressed and harvested as in (c). Soluble sugar levels (glucose, fructose, sucrose) are shown as mean ± SD from 3 biological replicates. (b-d) Different letters indicate significant differences at P < 0.05 (one-way ANOVA, Tukey’s HSD Test).

**Table 1.**
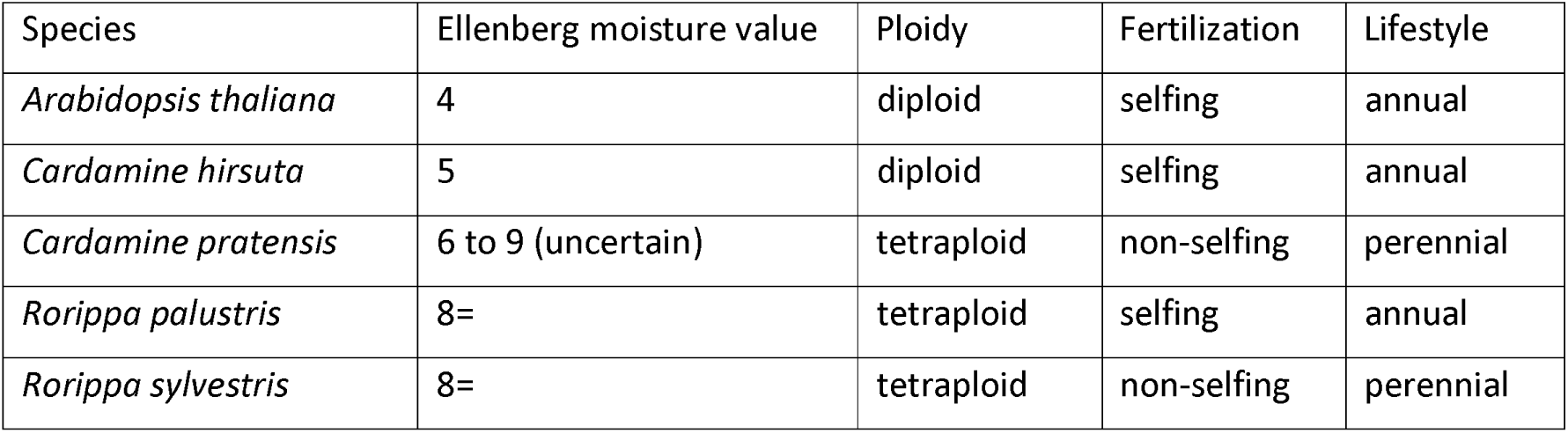
Ellenberg moisture values and characteristics of the studied species.

Next, we determined the underwater growth strategy in the five species by monitoring the youngest leaf during submergence. All five species lacked an escape response. Four species, *A. thaliana*, *C. hirsuta*, *R. palustris* and *R. sylvestris* had a significantly reduced leaf growth rate underwater during the first three days of submergence, while *C. pratensis* had similar growth rates under control and submerged conditions (Figure 1B).

The growth data was supplemented with the analyses of several relevant flood-tolerance related biochemical parameters. Alcohol dehydrogenase (ADH) activity was measured as a proxy for fermentation ability in response to any hypoxia experienced underwater. Though illuminated submergence does not cause hypoxia in the shoot (Arabidopsis, Lee et al. 2011; rapeseed, Wittig et al. 2021; watercress, Müller et al. 2021), at night local and subcellular oxygen deficiency might occur in submerged leaves. The two sensitive species (*A. thaliana, C. hirsuta*) did not enhance ADH activity after 24 or 48 h of submergence (Figure 1C). In contrast, *C. pratensis* and both *Rorippa* species had significantly higher ADH activities after 24 or 48 h of submergence (Figure 1C)

Soluble carbohydrates are the major substrate for glycolysis, which is the key energy source under hypoxia. The soluble carbohydrate content decreased in all five species within 24 and 48 h of submergence, but with some differences between species (Figure 1D). The sugar levels in control plants of *C. pratensis* and *R. sylvestris*, the two perennial species, were significantly higher compared with the other three species. After 24 and 48 h, the sugar levels were very low in most species, except for *R. sylvestris*, which showed significantly higher sugar levels compared with other species even after 48 h of submergence (Figure 1D).

### Flooding-sensitive Brassicaceae species have slower transcriptomic reconfiguration upon submergence

Biochemical and anatomical traits related to tolerance have a molecular origin. Furthermore, among tolerant species there is still considerable morphological variation, indicating a significant role for molecular tolerance. We therefore conducted transcriptome profiling to identify genes and key processes associated with flood tolerance, and potential novel tolerance regulators. To circumvent the lack of genome information for *C. pratensis*, *R. palustris* and *R. sylvestris* we made *de novo*-assembled transcriptomes for these species (Supplemental Figure S3). For optimal read mapping for *C. hirsuta* we created a genome-guided assembly to use as a reference based on the published *C. hirsuta* genome (Gan et al. 2016). Alignment success for expression quantification ranged from 70 to 87 % between the species (Supplemental Figure S4).

For inter-species transcriptomic comparisons, it is crucial to accurately identify orthologs. Therefore, we performed an all-vs-all blast with representative transcripts for each gene of the assembled transcriptomes (*C. hirsuta*, *C. pratensis*, *R. palustris*, *R. sylvestris*) and the primary transcripts of *A. thaliana* (Cheng et al. 2017) and *R. islandica* (Brassicales Map Alignment Project, DOE-JGI, http://bmap.jgi.doe.gov/). By clustering the resulting graph network of related sequences, we identified discrete groups of orthologous sequences, known as orthogroups (Emms and Kelly 2019, Supplemental data S1). We optimized the strength of grouping such that we maximized the number of orthogroups that cover all species (Supplemental Figure S5). This resulted in the identification of 16,902 universal orthogroups that were represented by 5 or 6 species (Figure 2A). For the species with available genome information (*A. thaliana*, *C. hirsuta* and *R. islandica*) relatively few genes could not be placed in a group of sequences from other species, compared to the *de novo* transcriptome-based assemblies (Figure 2B). Most orthogroups were represented by no gene or only one gene per species. However, the tetraploid species (*C. pratensis*, *R. sylvestris*, *R. palustris*) were frequently (∼35 %) present with two genes in an orthogroup (Figure 2C). When considering only the universal orthogroups, the vast majority of these orthogroups contained 1 gene for the diploid species (*C. hirsuta*, *A. thaliana*, *R. islandica*) and 2 genes for the tetraploid species (*C. pratensis*, *R. palustris*, *R. sylvestris*) (Figure 2D), which indicated good separation of homologous sequences into the smallest viable orthogroups containing most species.

**Figure 2.**
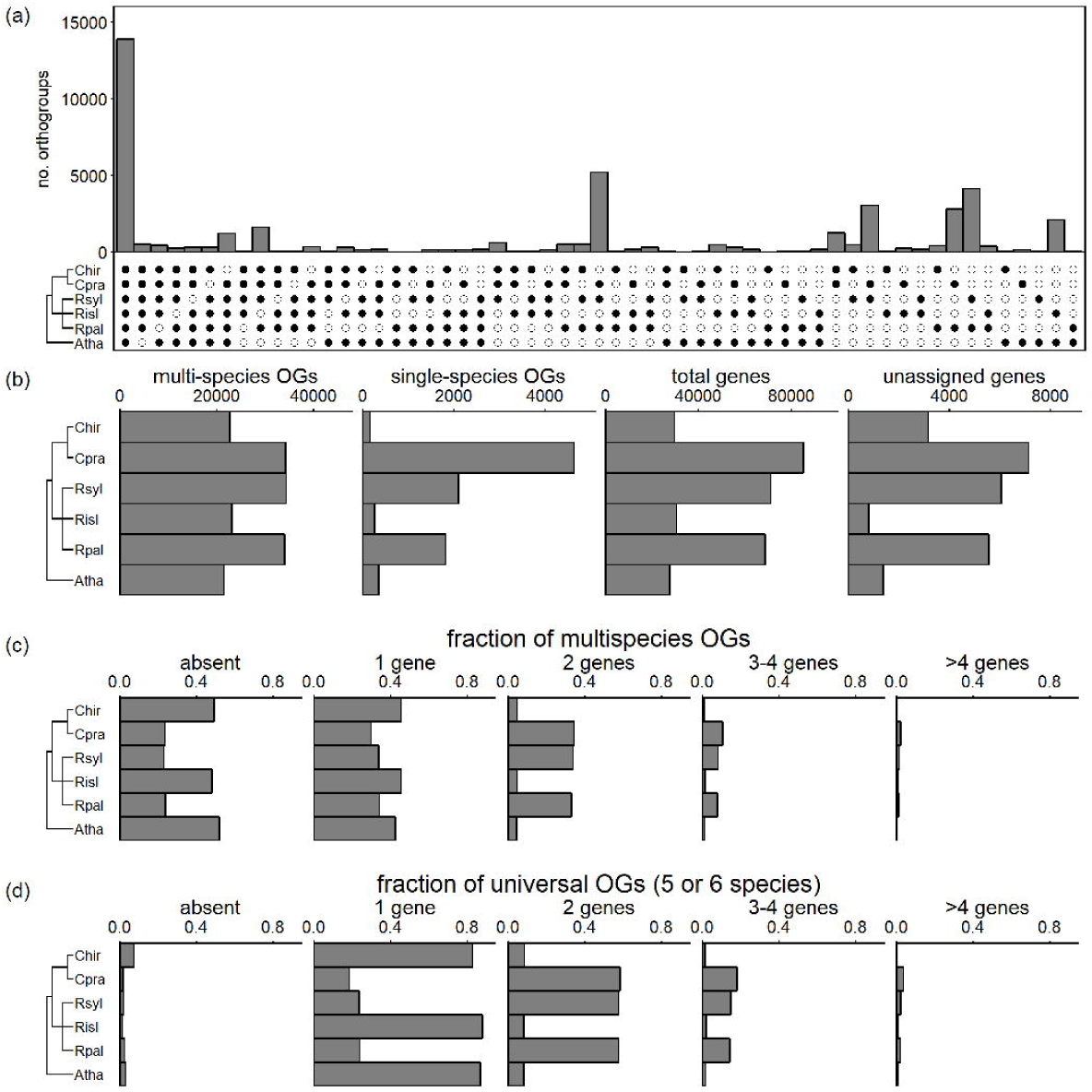
Identification of groups of orthologous sequence, orthogroups (OGs). (a) The number of orthogroups identified with representative sequences of the species indicated by filled circles. (b) Summary statistics of the genomic scope of the species considered for orthology analysis. Here multi-species OGs include sequences from at least two species; single-species OGs include two or more sequence from a single species that are grouped together because of high homology; total genes is the number of sequences on which orthology analysis was performed. An unassigned gene had insufficient homology to other sequences to be placed in a group with other sequences and so forms an OG by itself. (c) The proportion of Orthogroups that contain zero to many sequences of a particular species. (d) As (c), but then only considering Orthogroups that have sequences from at least five species. Chir – *C. hirsuta*, Cpra – *C. pratensis*, Rsyl – *R. sylvestris*, Rpal – *R. palustris*, Atha – *A. thaliana*.

To compare the transcriptomic reprogramming of the five species, we summarized orthogroup expression as the sum of all included transcripts as done previously for closely related species (Bräutigam et al. 2011, Van Veen et al. 2013). Overall, orthogroup expression was a reliable predictor of the expression level of the individual genes within that orthogroup, where for the separate species the estimated fold changes of the orthogroups explained 79 to 89 % of the variation found for individual genes (Supplemental Figure S6).

To check whether there were differences in the statistical strength to detect differential expression we compared the fold changes of differentially expressed genes to their overall sequencing depth. This showed a similar power to detect differential expression across species and timepoints (Supplemental Figure S7). Per species we fitted a full factorial model to retrieve the submergence response after 24 h, 48 h and an interaction effect to highlight temporal effects (Figure 3A). Additionally, we estimated the mean response to submergence over both timepoints by the main submergence effect in an additive model (Supplemental data S2, S3). Overall, the transcriptomic reconfiguration did not differ drastically between the five species. The number of differentially expressed orthogroups (DEOGs) after 24 or 48 h ranged from around 500 to 1,600 across all species (Figure 3A). The small number of orthogroups with a significant time*treatment interaction effect (Figure 3A) and a strong correlation between the timepoints (Figure 3B) indicated highly similar responses on both days. However, the more flood-sensitive *A. thaliana* and *C. hirsuta* had a delayed response in the downregulation of a considerable portion of their negatively regulated transcriptome.

**Figure 3.**
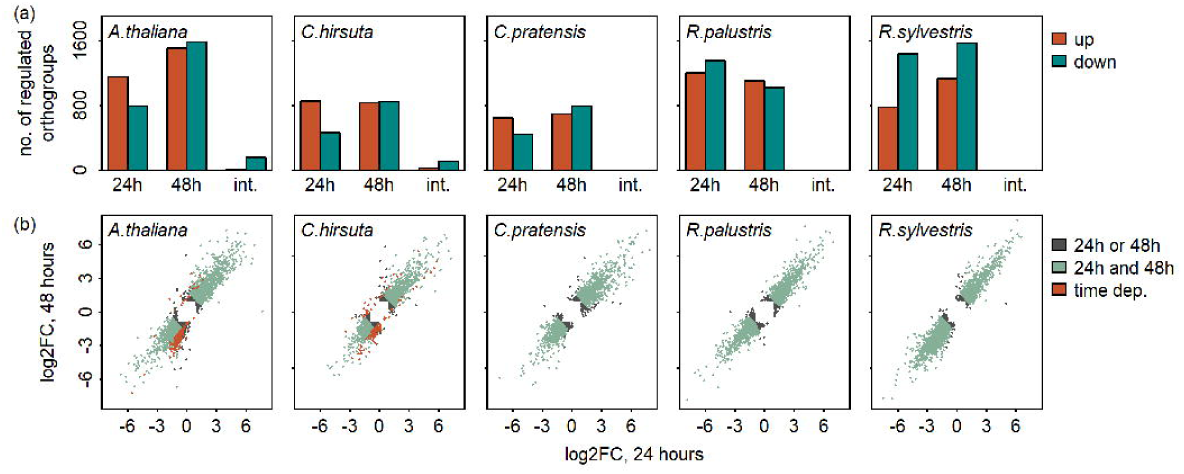
(a) The number of differentially expressed orthogroups (DEOG) after 24 and 48 h of submergence in the light compared to control after similar duration under aerated conditions, and whether the response at 48 h differs from the response at 24 h (time*treatment interaction: int.). Differential expression concerns a P_adj._ < 0.01 and a |log_2_FC| > 1. (b) Scatterplot depicting the relationship between transcriptomic differences at 24 and 48 h. Only orthogroups that are differentially expressed (24h, 48h or int.) are shown.

To functionally characterize the general transcriptomic response, a GO-term enrichment analysis was performed on the DEOGs per species (Supplemental data S4). This revealed among induced genes many GO terms associated with stress response (e.g, response to stimulus, response to stress, response to abiotic stimulus, response to light) and GO terms expected to be enriched under submergence, such as “response to hypoxia” and “response to ethylene”. The downregulated genes had fewer commonalities in their function, mainly related to synthesis of secondary metabolites.

### Inter-species comparison of ortholog behaviour reveals phylogenetic- and tolerance-dependent variation in the severity of transcriptomic responses

The above qualitative assessment provides a global picture of the transcriptomic behaviour in the five species. To quantitatively compare and pinpoint key sets of orthologous genes that can explain variation in flood tolerance or represent a conserved set of transcriptomic responses, we leveraged the established orthology (Figure 2) to directly contrast transcriptomic reconfiguration between species.

Direct pairwise comparison of DEOGs between species revealed a correlation in submergence responses. However, given a model where the transcriptional response would be identical between pairs of species, there remained a fair amount of unexplained variation with an average R^2^ value of 0.25 and extremes of 0.07 and 0.55 (Supplemental Figures S8, S9). These pairwise comparisons showed specific biases that indicated differences in the strength of up- or downregulation when submerged. To systematically portray these biases, we analysed the per-species deviation from the average submergence response across all five species (Figure 4). A sliding window across this mean response allowed us to assess biases (weaker or stronger than the mean response) in the strength of transcriptome reconfiguration. Regarding downregulation, for the tolerant *R. palustris* and *R. sylvestris* significantly more orthogroups showed a stronger downregulation after 24 h, with *R. sylvestris* sustaining this trend even after 48 h. In contrast, the downregulation of orthogroups in *C. hirsuta* and *C. pratensis* was weak, both at 24 and 48 h. *A. thaliana* over the whole had an average response after 24 h, but after 48 h downregulated genes had a stronger suppression and upregulated genes a stronger induction than the average of the species we assessed. Overall, this analysis combined with the number of observed DEOGs indicates that when genes are downregulated in the tolerant *Rorippa* species, the magnitude of regulation is strong. In comparison, the *Cardamine* species tend towards milder gene expression downregulation.

**Figure 4.**
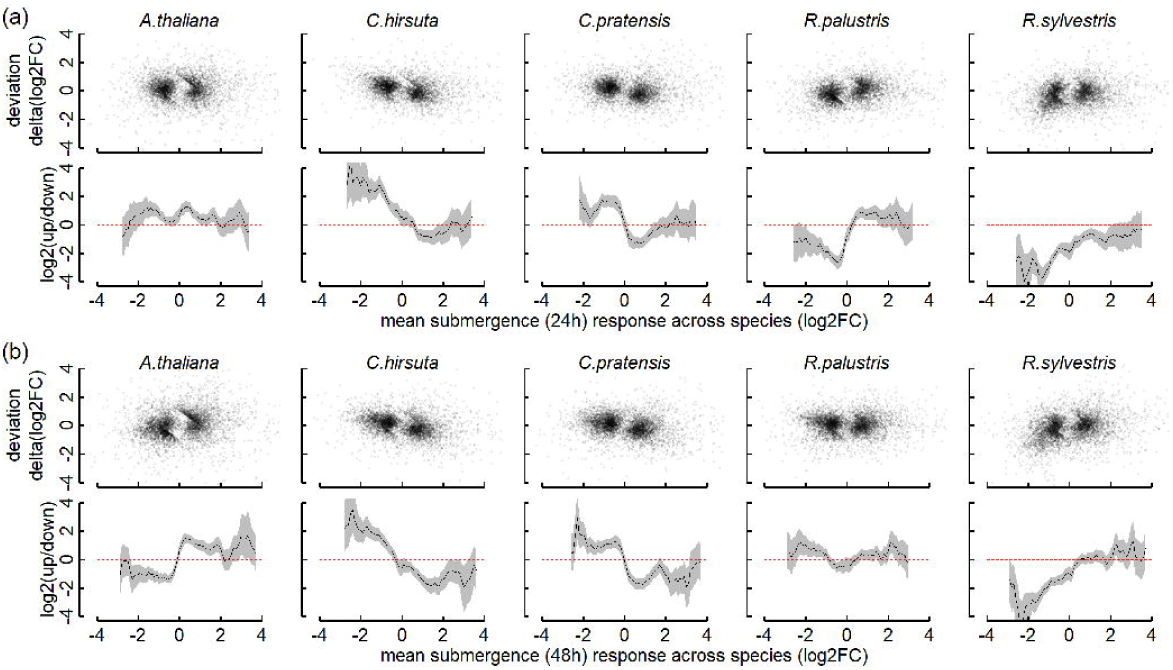
The response of individual species compared to average response across all species at 24 h (a) and 48 h (b). The deviation from the mean response considers the absolute distance of the log2FCs, where the red dashed line indicates a transcriptomic response identical to the species average (x-axis). The Log2 of the ratio of the number of positively and negatively deviating orthogroups (|delta(log2FC)| > 0.5) captures the bias of that species at any mean submergence response in a sliding window (width=0.5). The grey band considers the 95 % confidence interval (Exact Binomial Test), and the dashed red line indicates no deviation from the average. Orthogroups with P_adj._ < 0.001 and |log_2_FC| > 1 in at least 1 species were considered for the analysis.

To zoom in further on specific contrasting regulatory patterns between the species or highly conserved responses, we searched for orthogroups that showed significant regulation dependent on the species, genus or tribe by testing for a significant interaction of the phylogeny with treatment at either timepoint in three separate models (Figure 5, Supplemental data S5). Most between-species variation in transcriptomic reconfiguration, based on the number of significant orthogroups, was explained by individual species (species*treatment interaction; Figure 5A), with a vast amount being retained when species were nested into their corresponding genus. However, by nesting all Cardamineae species, a relatively poor phylogenetic signature was found based on the tribe*treatment interaction. This indicates that at least between the three genera studied here the flood responses are quite divergent. This reiterates the observation (Supplemental Figures S8, S9) that the genera were diverse in their submergence responses.

**Figure 5.**
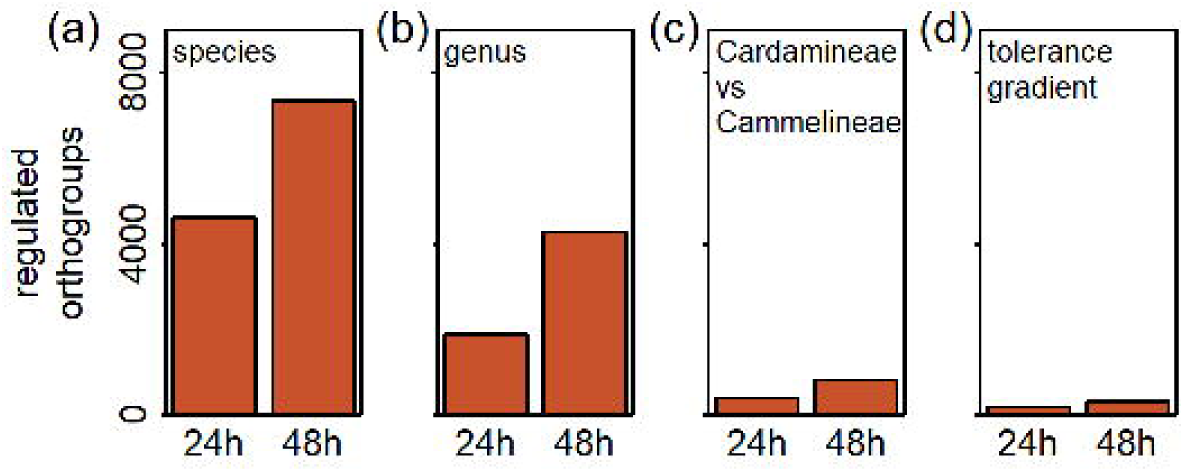
(a-c) The number orthogroups whose regulation is dependent on either the species (a), the genus (b) or the tribe (c) (species*treatment, genus*treatment, tribe*treatment; P_adj._ < 0.001). (d) The number of orthogroups whose magnitude of regulation only increases or decreases over successive tolerance groupings. Here *A. thaliana* is sensitive, *C. hirsuta* is intermediate, and *C. pratensis*, *R. palustris* and *R. sylvestris* are tolerant. In addition to an ordered change in magnitude of regulation these OGs require a significant effect of the grouping (P_adj._ <- 0.001) and considerable range in the magnitude of regulation|Log_2_FC_tolerant vs sensitive_| > 1).

Furthermore, we tested which orthogroups were significantly associated with the identified tolerance groups (Figure 1) and were regulated in line with a tolerance gradient from the sensitive *A. thaliana*, the moderately tolerant *C. hirsuta*, to the extremely tolerant *C. pratensis*, *R. palustris* and *R. sylvestris* (Figure 5D). Very few orthogroups had an expression profile that significantly matched the tolerance gradient (Rsyl, Rpal & Cprat ≥ Chir ≥ Atha). The tolerant Cardamineae tribe is not only a phylogenetic factor, by also provides a tolerance signature to further expand the candidate tolerance orthogroups.

The above test provided a platform to identify the highly conserved responses, transcriptomic adjustments with a phylogenetic signature, or those orthogroups with an expression associated with the observed tolerance gradient. First, to highlight the truly conservatively regulated genes that do not vary between the species we clustered the orthogroups without a species-specific effect (P_species*treatment_ > 0.05, P_treatment, mean across species_ < 0.01 and |log_2_FC _mean across species_| > 0.5). These 1,095 DEOGs were split into four distinct clusters (Figure 6, Supplemental data S6) where the majority of orthogroups showed a mild up- or downregulation upon submergence (cluster 3, 515 DEOGs; cluster 1, 414 DEOGs), and 53 orthogroups that were strongly upregulated (cluster 4) or 113 orthogroups with moderate downregulation (cluster 2). GO categories that were enriched among the mildly induced orthogroups included biotic defence responses, indole glucosinolate metabolism and autophagy (Supplemental data S7). GO enrichment among mildly downregulated orthogroups were concerned mostly with mitochondrial respiration and electron transport chain, and possibly cell division via spindle formation. The strongly induced or downregulated clusters had only poor GO enrichment, partly indicative of the smaller cluster size, but also a functionally diverse set of orthogroups. However, the strongly downregulated orthogroups were enriched in xylan metabolism and leaf development.

**Figure 6.**
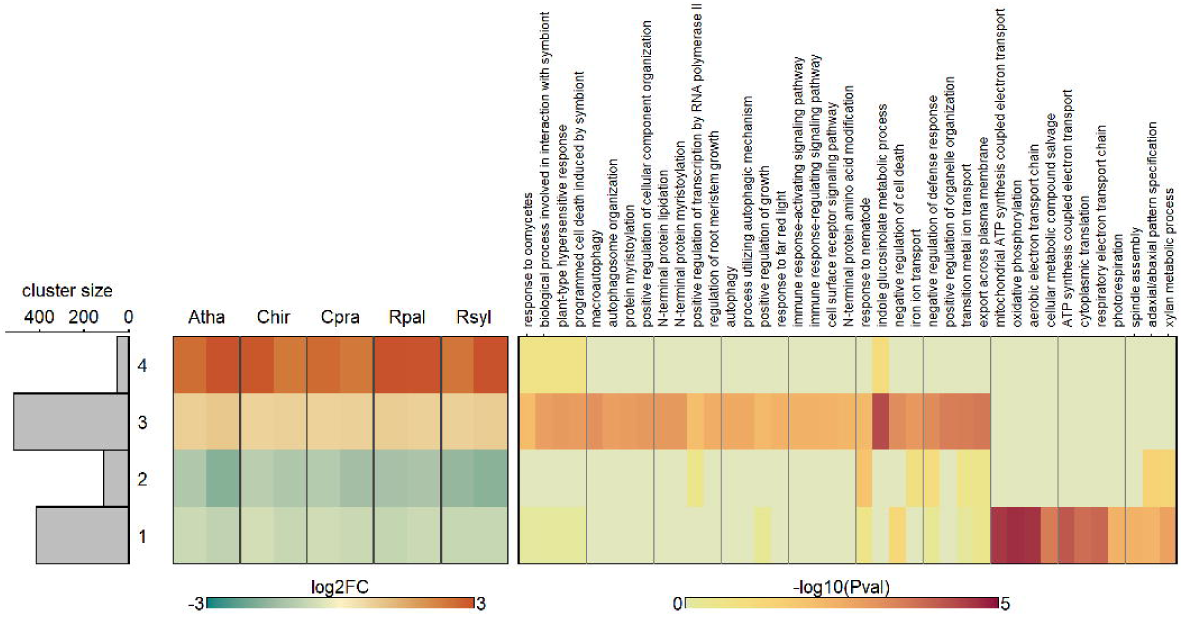
Clustering of the orthogroups without any species effect. For these no significant effect of species (P_adj._ > 0.05; Figure 5A), but a significant mean response across all species (P_adj._ < 0.01 and |log_2_FC| > 0.5) was observed. The data was hierarchically clustered based on Euclidean distances and an agglomeration by a minimum variance method. Only GOterms with at least 80 OGs, 5 DEOGs and P < 0.01 are shown. For brevity only Biological Process GOterms are depicted. Cluster assignment of individual OGs and complete GO enrichment analysis can be found in Supplemental data S6 and S7.

To highlight any phylogenetic signatures in the transcriptomic reconfiguration, we also highlighted the genus-dependent flood responses (P_genus*treatment_ < 0.001 and |log_2_FC_between any genus_ | > 1, Figure 7A). We found genus to be the lowest phylogenetic level that still retained a strong signature of evolutionary divergence in submergence responses (Figure 5). Clustering and GO term analysis of these 2,609 orthogroups revealed strong enrichment of cell cycle, DNA replication, disaccharide metabolism related terms among orthogroups that were downregulated particularly in the *Rorippa* species (Cluster 7, 233 DEOGs, Supplemental data S6, S7). Regarding orthogroups especially induced in *Arabidopsis*, chlorophyll breakdown-related terms were enriched (Cluster 6, 246 DEOGs). The *Cardamine* genus also showed unique patterns of induction or suppression. Here anthocyanin metabolism and responses to cytokinin were enriched among orthogroups that were mildly downregulated compared to *Rorippa* and *Arabidopsis* (Cluster 8, 46 DEOGs). Also, the *Cardamine* species had sets of orthogroups that were only poorly upregulated compared to the other species (Cluster 3, 74 DEOGs), or were only upregulated in the *Cardamine* (Cluster 4, 115). However, there was no clear enrichment of GO terms among these clusters.

**Figure 7.**
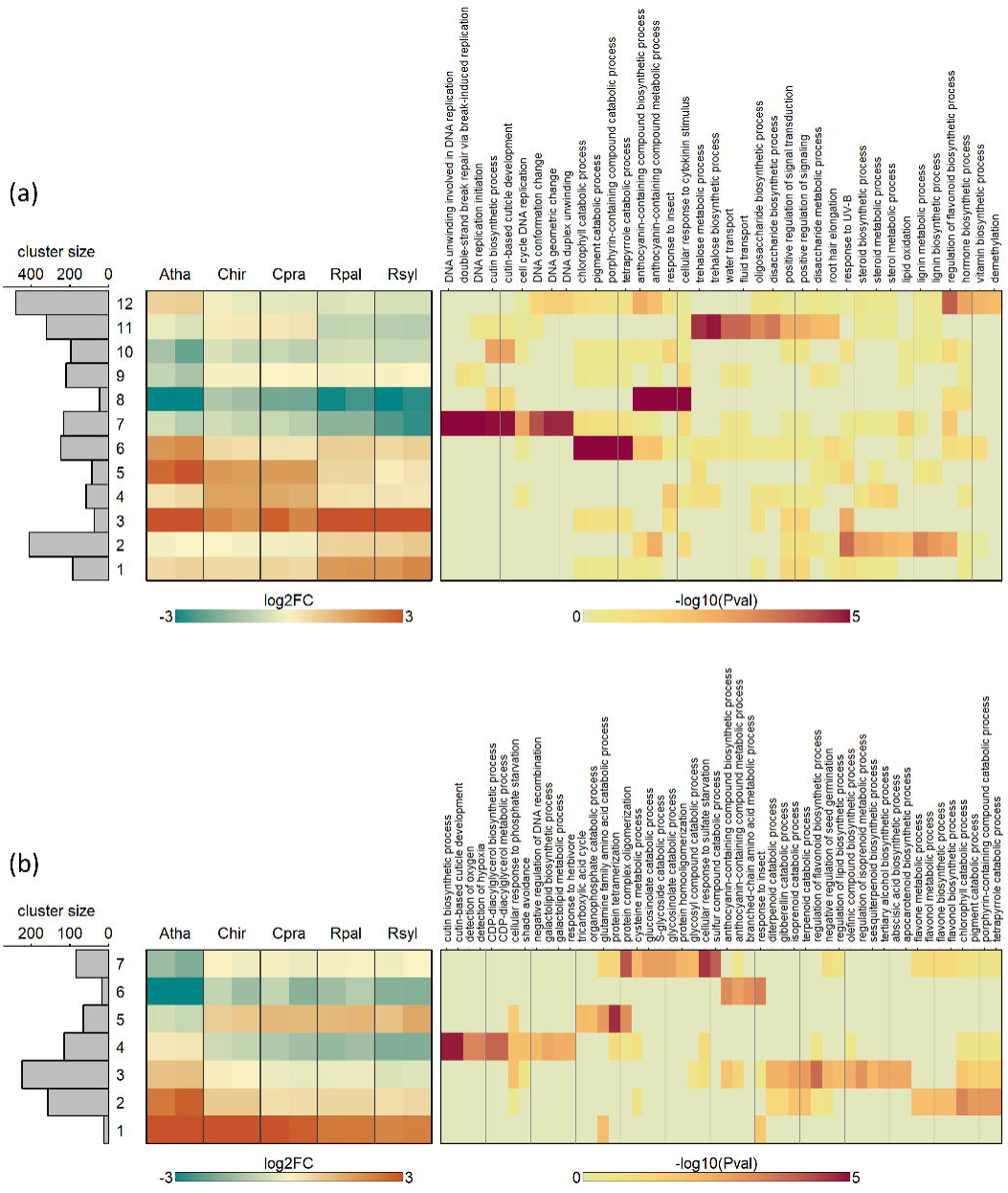
Clustering of the orthogroups with divergent transcriptional response between species. (a) Clustering and GO enrichment of OGs whose transcriptional response depends on the genus, thus OGs where a P_adj._ < 0.001 (Figure 5B) and |log_2_FC_between any genus_| > 1 was observed. In (b) transcriptional responses associated with tolerance are clustered. These are the OGs where the transcriptional response depends on the tribe, thus with a P_adj._ < 0.001 (Figure 5C) and a |log_2_FC_Cardimaneae vs A.thaliana_ | > 1), and also those OGs that followed the tolerance gradient (Figure 5D). The data were hierarchically clustered based on Euclidean distances and an agglomeration by a minimum variance method. Only GOterms with at least 80 OGs, 5 DEOGs and P < 0.01 are shown. For brevity only Biological Process GOterms are depicted. Cluster assignment of individual OGs and complete GO enrichment analysis can be found in Supplemental data S6 and S7.

Relatively few genus-specific orthogroups showed a pattern that would support a unified approach to achieve tolerance, i.e. up or down in all tolerant species. However, we ultimately sought to identify the specific molecular components that could be instrumental in conferring submergence tolerance or sensitivity among the five selected species. We thus selected the orthogroups that had a tribe-specific effect, since all Cardamineae are at least moderately tolerant (Figure 1, P_tribe*treatment_ < 0.001 and |log_2_FC_tribe*treatment_| > 1). And we selected those with a regulatory pattern that follows the tolerance gradient (Figure 1, P_tol*treatment_ < 0.001 & *Rsyl*, *Rpal* & *Cpra* ≥ *Chir* ≥ *Atha*). The resulting 675 orthogroups were placed into seven clusters of similar regulation patterns (Figure 7B, Supplemental data S6, S7), which showed that the tolerant species mostly lacked or had a reduced up- and downregulation compared to the sensitive *A. thaliana* and moderately tolerant *C. hirsuta*. Among these upregulated orthogroups several GO terms associated with chlorophyll breakdown (Cluster 2, 157 DEOGs) and secondary metabolism were enriched (cluster 3, 224 DEOGs), whereas among the downregulated orthogroups enrichment was in terms associated with anthocyanin and glucosinolate metabolism (Cluster 7, 84 DEOGs). The orthogroups that did show a stronger regulation, either up or down, in the tolerant species had GO enrichment for terms associated with protein complex oligomerization and the TCA cycle for upregulated orthogroups (Cluster 5, 66 DEOGs). Downregulated orthogroups were overrepresented by cutin biosynthesis-related terms, shade avoidance and oxygen perception (Cluster 4, 115 DEOGs).

### Hypoxic responses are minor, whilst carbon starvation and ethylene-regulated transcripts are the most responsive in the submergence transcriptome

Hypoxia, ethylene and carbon starvation are strongly associated with submergence. Indeed, these aspects surfaced in the unbiased transcriptome analysis and phenotypic characterization of the five species. Given their importance, we also investigated transcripts associated with these processes in a targeted approach.

Mustroph et al. (2009) identified a subset of genes induced upon hypoxia regardless of tissue and cell type in *A. thaliana*, which are widely considered a core set of hypoxia responsive genes (HRGs). HRGs responded similarly in the five tested species in response to submergence (Figure 8A, Supplemental data S8), but were not consistently upregulated. Only *HUP54*, *ACHT5*, *PP2-A13*, and an unknown protein were strongly induced. The HRGs typically considered as highly reliable hypoxia-response indicators, namely *PCO1/2*, *LBD41*, *ADH1* (Gasch et al. 2016, Weits et al. 2014), were not induced. Comparison of microarray data across a variety of hypoxia studies in *A. thaliana* and other species has yielded a broader set of genes that are conserved in their upregulation upon hypoxia (Mustroph et al. 2010). Many of these genes were not induced upon submergence (Figure 8B, Supplemental data S8). The weak hypoxic signature in this study was expected since illuminated shoots rarely experience hypoxia (Colmer & Pedersen 2008, Lee et al. 2011, Vashisht et al. 2011, van Veen et al. 2013, Müller et al. 2021, Wittig et al. 2021).

**Figure 8.**
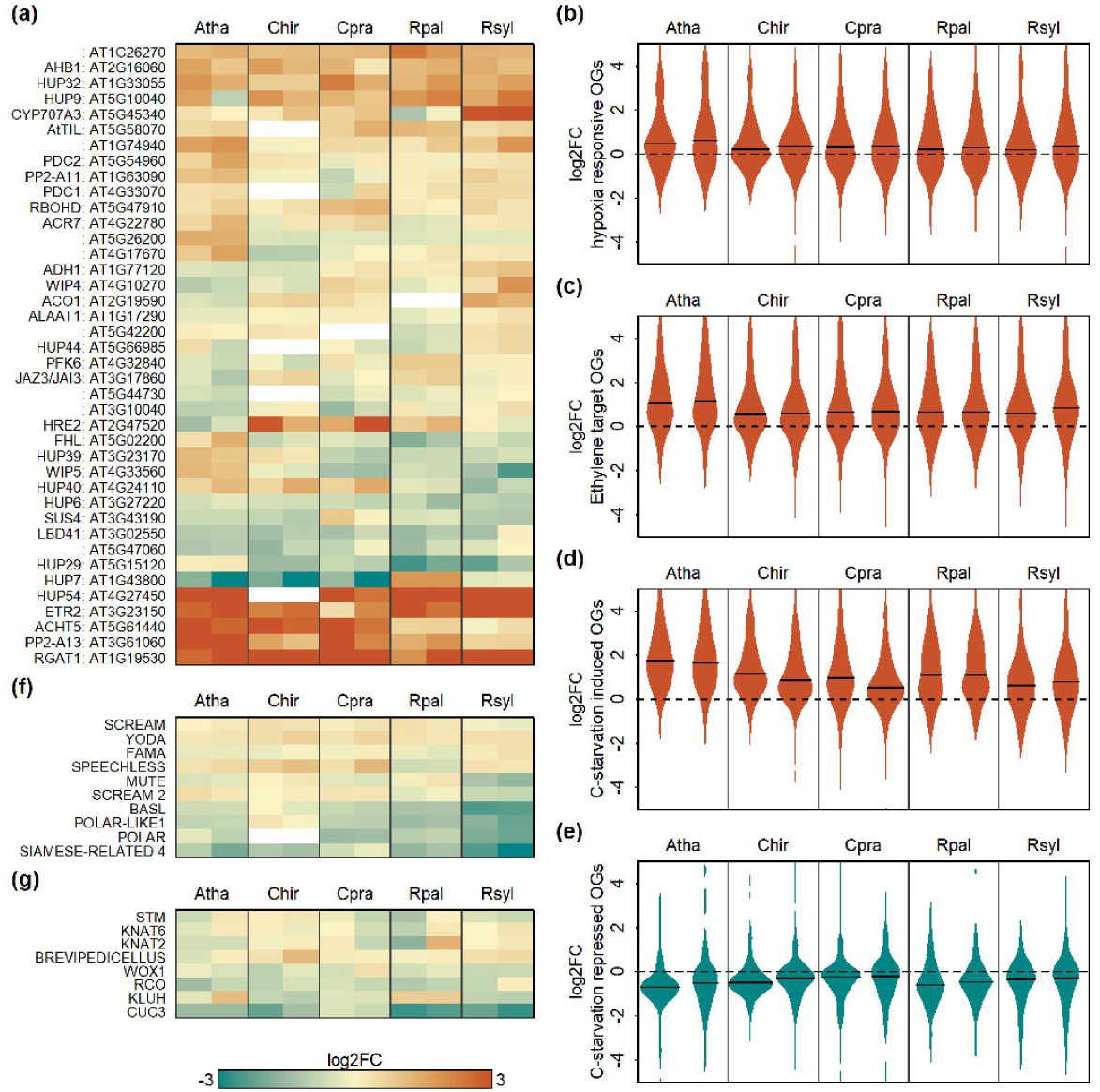
Targeted analysis of key flood adaptive responses. (a) transcriptional response of core hypoxia responsive genes (HRGs), which are hypoxia responsive regardless of cell type, as defined by Mustroph et al. (2009). (b) Density of the response of conserved hypoxia responsive genes that are regulated across a wide range of plant species and hypoxia studies, as defined by Mustroph et al. (2010). (c) Density of ethylene-responsive orthogroups, which in *A. thaliana* contain an EIN3 binding domain in their promotor and are induced by ethylene, as defined by Chang et al. (2013). (d) Density of carbon starvation-induced orthogroups as defined by Usadel et al. (2008) by combination of sugar feeding and low CO_2_ studies. (e) is as (d), but with carbon starvation-repressed orthogroups. The horizontal line in (b), (c), (d) and (e) indicates the median log2FC of the set of orthogroups. (f) Heatmap of key players of stomata development. Induction of these genes is associated with more stomates (Smit and Bergmann 2023). (g) Heatmap of key leaf developmental players required for leaf dissection (Bhatia et al. 2021).

The gaseous hormone ethylene, rather than hypoxia, is considered a more reliable cue for submergence (Sasidharan et al. 2018). Indeed, the GOterm “Response to ethylene” was significantly enriched (Pval: from 5.4E-10 to 1.3E-19) among upregulated genes in all species at both timepoints (Supplemental data S4). A focused investigation of the response of DEOGs considered to be direct targets of ethylene signalling by virtue of both an EIN3 binding domain in their promotor and induction upon ethylene treatment in *A. thaliana* (Chang et al. 2013) confirmed the presence of an ethylene signature upon submergence in our data (Figure 8C, Supplemental data S8). Of all species, *A. thaliana* had the strongest ethylene response, which was reflected in the highest median log2FC. However, this could also be an artifact of the *A. thaliana*-biased selection of ethylene target orthogroups.

Though tolerant species maintained a better sugar status than sensitive species, we observed a drop in sugar levels and an increase in autophagy-related transcripts across all five species (Figures 1D, 6). Carbon starvation followed by sugar feeding of seedlings and low CO_2_ treatment of illuminated rosettes provided a reliable set of carbon responsive genes in *A. thaliana* (Usadel et al. 2008). Taken together, these starvation-induced and -repressed genes in the five species studied here showed that all species had transcriptome reconfiguration associated with carbon starvation, and that this reconfiguration was the strongest in *Arabidopsis* (Figure 8D-E).

Hampered photosynthesis underwater, leading to carbon limitation could explain the dominant carbon starvation effects in the transcriptomes. CO_2_ limitation is specifically reflected by the induction of alanine:glyoxylate aminotransferase, a key player in photorespiration, in all species apart from *C. pratensis* (OG0004610 & OG0004408, Supplemental data S3, Liepman & Olsen 2003, Niessen et al. 2012). Several frequently flooded plant species have evolved mechanisms to mitigate diffusion limitations. These encompass a reduction or absence of both the cuticle and stomatal abundance, and higher specific leaf area (Van Veen & Sasidharan 2021). Since cuticle biosynthesis had the strongest downregulation in the *Rorippa* species (Figure 7), we explored the possibility of further morphological adaptations that improve underwater photosynthesis, and also explain differences in carbon status. Indeed, key players of stomatal development, such as *SCREAM2*, *BASL*, *POLAR*, *POLAR LIKE* and *SIAMESE RELATED4* (Smit & Bergmann, 2023), were downregulated particularly in the tolerant species (Figure 8F). *R. aquatica* acclimates to underwater conditions by increasing leaf dissection (Nakayama et al. 2014). However, the known key players in the control of leaf dissection (Bhatia et al. 2021) showed no clear pattern in transcriptional reconfiguration (Figure 8G).

## Discussion

### The Cardamineae provide a valuable resource for studying flood-tolerance traits

Several species from the Brassicaceae family have been investigated for flood resilience. Notably, the Cardamineae tribe contains several highly flood-tolerant species, including various members of the *Rorippa* genus (Akman et al. 2012, Van Veen et al. 2014, Nakayama et al. 2014) and *Nasturtium officinale* (Müller et al. 2021). These studies have yielded valuable insights into the adaptive strategies of these species. For example, regulation of the shoot escape strategy in *N. officinale* (Müller et al. 2021) bypassed the conserved ethylene-ABA-GA hormonal network. The formation of distinct aquatic and terrestrial leaves in *R. aquatica* has made it a model to study environmental control of leaf development (Nakayama et al. 2014).

Here we leveraged the diverse range of flood tolerance exhibited within the Brassicaceae to study molecular responses to submergence. We extended the species palette by incorporating two *Cardamine* species and found a medium to high tolerance. By also including the sensitive *A. thaliana*, our panel represented a broad tolerance range across five species. All five species displayed a quiescent strategy underwater with submergence tolerance spanning 4 to >10 weeks. This wide survivability range surpasses the natural variation in submergence tolerance observed among natural Arabidopsis accessions (Vashisht et al. 2011). The selection of these quiescent species therefore provides a robust study system to explore flooding-tolerance mechanisms in plant species enabling the capture of adaptive signatures at a much broader genetic scale.

A notable constraint in comparisons of molecular stress responses of a wider palette of species is the capacity to compare transcriptomes effectively and accurately. In the early days, these were partly related to capacity limitations and high costs of NGS technologies, with most studies confined to a single or pair of contrasting species to explore environmental responses. Examples regarding flooding include escape and quiescence responses in the genus *Rumex* (Van Veen et al. 2013), aquatic leaves in waterlily (Wu et al. 2017), and heterophylly in *Ranunculus trichophyllus* (Kim et al. 2018) and *Rorippa aquatica* (Ikematsu et al. 2023).

Some studies resorted to the use of Arabidopsis microarrays, but the requirement for high sequence similarity restricted species choice, e.g *Rorippa* (Sasidharan et al. 2013) and *Cardamine* (Shimizu-Inatsugi et al. 2017). The advent of affordable, high-resolution NGS platforms has reduced such challenges and permits a more diverse choice of plant species driven by scientific questions rather than technological constraints. However, gene diversification among distantly related plant species still poses challenges for direct ortholog comparisons. For example, a comparative study on the gene regulatory networks of hypoxia signalling ranging from monocots to dicots was restricted to less than 7,000 shared gene families (Reynoso et al. 2019). This loss of resolution restricts findings to highly conserved genes and conserved responses. However, a study on fungal responses in the core eudicots distinguished < 8,000 core gene families but managed to leverage the diversification in the gene universe to explain adaptation by regulation specifically in lineage-specific gene families (Sucher et al. 2020).

In contrast, our study over a relatively narrow phylogenetic distance identified 16,902 universal orthogroups. At the same time, we identified a substantial diversity in stress responses. Orthologs between species were identified mostly on a 1-to-1 basis or 1-to-2 or 2-to-2 basis in the case of tetraploid species. This permitted disentanglement of responses at a high resolution and direct ortholog level comparison and clustering. This study thus goes beyond describing processes, tracing the actual genes that evolved to mediate stress adaption. The transcriptomic signatures suggest different routes towards tolerance with different identified gene sets in *Rorippa* and *Cardamine*. Also, a direct ortholog-based comparison with sharply separated gene relationships provided the scope to do a quantitative assessment of flood responses, i.e. mild vs strong upregulation, rather than a qualitative resolution where gene families are classified as up, down, or neither. A quantitative analysis was instrumental in identifying the attenuated regulation in *Cardamine* vs the strong downregulation in *Rorippa*, which would not have readily emerged out of a qualitative transcriptome assessment. The Cardamineae tribe thus provides an excellent study system to disentangle flood adaptions and tolerance in greater detail, for which we here provide the first step.

### Carbon starvation, autophagy and biotic defences are conserved responses to submergence

Despite considerable variation in flood tolerance, several processes were induced and suppressed equally across the five species. Of note were the activation of pathways associated with biotic stress and autophagy (Figure 6). Upregulation of the defence response during flooding has been observed in several transcriptomic studies encompassing Brassicaceae species, but also in *Rumex* (Müller et al. 2021, Wittig et al. 2021, Van Veen et al. 2013, Van Veen et al. 2016). The reason for this conserved induction of defence response remains unresolved but is potentially associated with ethylene accumulation in submerged tissues. Ethylene is an established participant in the signalling network regulating plant defence responses (Broekgaarden et al. 2015). Alternatively, defence signalling might be activated to combat elevated infection pressure following submergence (Hsu et al. 2013).

Autophagy is required to maintain cellular energy homeostasis when the substrate demand for glycolysis or respiration exceeds supply, which typically occurs during flooding (Dalle Carbonare et al. 2023). The importance of autophagy for flood tolerance is apparent from the increased flood sensitivity observed in various autophagy mutants (Chen et al. 2015). General stress-responsive GO terms and carbohydrate starvation were among the common responses (Supplemental data S4) and have been identified in a variety of tolerant (Müller et al. 2021, van Veen et al. 2013) and sensitive species (Wittig et al. 2021, van Veen et al. 2016). Focused analysis of this starvation signature indicated a more attenuated response in species from the tolerant Cardamineae tribe (Fig. 8D). For the *Rorippa* genus this matched the better maintenance of sugar levels, but this was not the case for the *Cardamine* species (Figure 1D). Sensitive rapeseed plants also had very low carbohydrate levels already after 3 h of submergence in light, with levels not rising again within 24 h of submergence (Wittig et al. 2021). The underlying causes of the strength of starvation signalling might lie in differences in underwater photosynthesis, carbon utilization rates, or sensitivity to starvation cues. Regardless, all species experienced carbon starvation.

Plant hypoxia responses are associated with the signature upregulation of a conserved gene set (Mustroph et al. 2009, 2010). In this study a limited number of these genes were induced (Figure 8A and B). Despite the likely absence of day-time hypoxia in our experimental system, the upregulation of these hypoxia-responsive genes could potentially be attributed to ethylene inducibility of some core hypoxic genes (Van Veen et al. 2013, Hartman et al. 2019). Illuminated submerged conditions do permit some photosynthesis, providing some energy and oxygen, which combats hypoxia. Light availability can greatly improve longevity underwater (Mommer et al. 2006, Vashisht et al. 2011). Despite a minor induction of *ADH* and *PDC* transcription (Figure 8A), we observed increased ADH activity, especially in tolerant species (Figure 1C), which might be retained from night-time hypoxia, possibly combined with minor day-time transcription driven by ethylene. These results show a positive association between the strength of hypoxic acclimation and flood tolerance. However, such relationships are not universally found (Dalle Carbonare et al. 2023). In contrast, starvation and defence responses seem more conserved transcriptomic responses to submergence.

### Phylogenetically distinct routes towards flood tolerance associate with transcriptional sensitivity and quiescence

There were many species-specific and genus-specific orthogroups, compared to conserved and tolerance-related responses (Figures 3A, 5). There was a strong phylogenetic signature, indicating distinct transcriptional reconfiguration for the *Cardamine* and *Rorippa* genus in comparison to *A. thaliana* to achieve flood tolerance. Observations of an aquatic habitat among angiosperm phylogenies suggested that an aquatic/amphibious lifestyle evolved over 200 times (Cook 1999). Given the possibility of alternative tolerance strategies already within the narrow genetic range of the Cardamineae suggests that an amphibious lifestyle was adopted many more times.

Both *Cardamine* species had a weaker down- and upregulation, thereby maintaining a transcriptomic signature closer to that of an aerated plant. This is especially pronounced with regard to chlorophyll breakdown, compared to *Rorippa* and *Arabidopsis* (Figure 7). The advantage of a weak-response strategy is apparent from *A. thaliana* mutants retarded in senescence that perform better than wild type at non-lethal flood durations (Rankenberg et al. 2024). Similarly, an *A. thaliana* accession with superior flooding recovery had an overall weaker transcriptomic reconfiguration when flooded (Yeung et al. 2018). However, the transcriptome configuration both stems from the plant’s physiology, and affects the physiology itself. The stronger senescence signature found here for *A. thaliana* compared to the tolerant species (Figure 7) could result from greater stress caused by the low flood tolerance. However, the observation that delayed senescence caused by genetic or chemical ethylene signalling inhibition improves performance (Rankenberg et al. 2024), favours a gene regulatory program.

In contrast, an absence or a delay in transcriptomic reconfiguration has also been associated with heightened hypoxia sensitivity. Pre-treatment with ethylene resulted in faster hypoxic gene induction in *Rumex palustris* and *A. thaliana*, corresponding with improved anoxia tolerance (Van Veen et al. 2013, Hartman et al. 2019). Strikingly, the two most sensitive species, *A. thaliana* and *C. hirsuta*, were the slowest in their downregulation response.

Opposite to the *Cardamine* species, both *Rorippa* species displayed strong downregulation (Figure 4). The clustering and GO enrichment indicated that this was primarily related to cell-cycle activity (Figure 7). Although leaf length measurements indicated that *Rorippa* species were not more quiescent than *A. thaliana* (Figure 1B), reduced cell-cycle activity suggests that particularly the development of new or younger leaves would be affected. Notably, shade avoidance, which affects primarily cell elongation, was also downregulated especially in the tolerant species. Activating shade avoidance is a key mechanism of the escape response of *Rumex palustris* (Van Veen et al. 2013). Overall, the observations emphasize the importance of quiescence, which might encompass more than the phenotypically discernible. With strong cell-cycle suppression, the *Rorippa* species studied here might represent a prime example of successful quiescence. This would be an effective strategy to reduce carbon and energy expenditure and is reflected in the milder sugar starvation in the two *Rorippa* species.

Heterophylly is advantageous under prolonged illuminated submerged conditions, a trait found in *R. aquatica* (Nakayama et al. 2014). A transcriptome response associated with reduced cutin biosynthesis was one of the few universally adopted by tolerant Cardamineae species (Figure 7). Furthermore, the *Rorippa* species had strongly reduced stomatal development, at least at the transcriptional level (Figure 8). These observations might simply reflect the stalling of leaf production. Nonetheless, there is strong variation in the strength of heterophylly, e.g. *Rumex palustris* vs *Ranunculus trichophyllus* (Mommer et al. 2005, Kim et al. 2018), and even minor changes could improve underwater photosynthesis and tolerance.

## Conclusions

We show that a focus on a narrow phylogenetic range, but with numerous species and diverse phenotypic data, permits an analysis that goes beyond a qualitative inventory. The capacity to separate orthologs at a high resolution provided a platform to separate mild from strong transcriptomic reconfiguration, and to directly cluster orthologs between species. Consequently, we could separate specific genes, rather than processes, that would have changed their regulation resulting in flood tolerance. We suggest that limiting senescence contributes to tolerance. However, to maintain energy homeostasis under a negative carbon balance some senescence would be unavoidable. Strong cell-cycle suppression seems an important feature of *Rorippa* and key to maintaining energy homeostasis, at least in the early phase of submergence. Fast and strong hypoxia signalling is beneficial for hypoxia tolerance, but might not be relevant to illuminated submergence, especially here, where quiescence and limiting carbon seem imperative. Dissecting exactly which condition favours a fast, slow, weak or strong response will be a crucial challenge for future work.

## Acknowledgements

We thank Pia Schuster and Marina Selle for technical assistance. Andrew M. Latimer is acknowledged for his logistic support for RNA-seq library preparations.

## Supplemental Data

### Supplemental methods

Figure S1. Phylogenetic and phenotypic characterization of selected Brassicaceae

Figure S2. Pictures of plants after submergence stress and a two-week-recovery phase.

Figure S3. N50 to N90 statistics of all assembled transcripts and the *A. thaliana* transcriptome.

Figure S4. Percentage of reads that successfully mapped to the reference transcriptome.

Figure S5. Inflation parameter optimization in Orthofinder.

Figure S6. The extent to which the fold change of a transcript can be predicted by the fold change of the orthogroup.

Figure S7. The fold change of orthogroups compared to their mean sequencing depth across all samples.

Figure S8. Pairwise comparison of 24 h flood responses between all species combinations.

Figure S9. Pairwise comparison of 48 h flood responses between all species combinations.

Data S1: Assignment of orthogroups (.txt.gz)

Data S2a: Fold Changes by gene of the universal orthogroups (.txt.gz)

Data S2b: Fold Changes by gene of the non-universal orthogroups (.txt.gz)

Data S3A: Fold Changes of the universal orthogroups (.txt.gz)

Data S3B: Fold Changes of the non-universal orthogroups (.txt.gz)

Data S4A: GO enrichment of downregulated orthogroups (.txt.gz)

Data S4B: GO enrichment of upregulated orthogroups (.txt.gz)

Data S5: The effect of phylogeny and tolerance grouping on submergence (.txt.gz)

Data S6: Membership of orthogroups to clusters (.txt.gz)

Data S7A: GO enrichment among clusters with no species effect (.txt.gz)

Data S7B: GO enrichment among clusters with genus-specific responses (.txt.gz)

Data S7C: GO enrichment among clusters with tolerance-specific responses (.txt.gz)

Data S8: Orthogroups and fold changes of the targeted processes of interest (.txt.gz)

## References

Akiyama R., Sun J., Hatakeyama M., Lischer H.E.L., Briskine R.V., Hay A.,… Shimizu-Inatsugi R. (2021) Fine-scale empirical data on niche divergence and homeolog expression patterns in an allopolyploid and its diploid progenitor species. New Phytologist 229, 3587–3601.

Akman M., Bhikharie A.V., McLean E.H., Boonman A., Visser E.J.W., Schranz M.E. & Van Tienderen P.H. (2012) Wait or escape? Contrasting submergence tolerance strategies of *Rorippa amphibia*, *Rorippa sylvestris* and their hybrid. Annals of Botany 109, 1263– 1276.

Akman M., Bhikharie A.V., Mustroph A. & Sasidharan R. (2014) Extreme flooding tolerance in *Rorippa*. Plant Signaling & Behavior 9, e27847.

Bhatia N., Runions A. & Tsiantis M. (2021) Leaf Shape Diversity: From Genetic Modules to Computational Models. Annual Review of Plant Biology 72, 325–356.

Brassicales Map Alignment Project, DOE-JGI, http://bmap.jgi.doe.gov/

Bräutigam A., Kajala K., Wullenweber J., Sommer M., Gagneul D., Weber K.L.,… Weber A.P.M. (2011) An mRNA Blueprint for C4 Photosynthesis Derived from Comparative Transcriptomics of Closely Related C3 and C4 Species. Plant Physiology 155, 142–156.

Bray N.L., Pimentel H., Melsted P. & Pachter L. (2016) Near-optimal probabilistic RNA-seq quantification. Nature Biotechnology 34, 525–527.

Broekgaarden C., Caarls L., Vos I.A., Pieterse C.M.J. & Van Wees S.C.M. (2015) Ethylene: traffic controller on hormonal crossroads to defense. Plant Physiology, pp.01020.2015.

Camacho C., Coulouris G., Avagyan V., Ma N., Papadopoulos J., Bealer K. & Madden T.L. (2009) BLAST+: architecture and applications. BMC Bioinformatics 10, 421.

Carlson M (2019a) org.At.tair.db: Genome wide annotation for Arabidopsis. R package version 3.8.2.

Carlson M (2019b) GO.db: A set of annotation maps describing the entire Gene Ontology. R package version 3.8.2.

Chang K.N., Zhong S., Weirauch M.T., Hon G., Pelizzola M., Li H.,… Ecker J.R. (2013) Temporal transcriptional response to ethylene gas drives growth hormone cross-regulation in Arabidopsis. eLife 2, e00675.

Chen L., Liao B., Qi H., Xie L.-J., Huang L., Tan W.-J.,… Xiao S. (2015) Autophagy contributes to regulation of the hypoxia response during submergence in *Arabidopsis thaliana*. Autophagy 11, 2233–2246.

Cheng C., Krishnakumar V., Chan A.P., Thibaud-Nissen F., Schobel S. & Town C.D. (2017) Araport11: a complete reannotation of the *Arabidopsis thaliana* reference genome. The Plant Journal 89, 789–804.

Colmer T.D. & Pedersen O. (2008) Oxygen dynamics in submerged rice (*Oryza sativa*). New Phytologist 178, 326–334.

Combs-Giroir R. & Gschwend A.R. (2024) Physical and molecular responses to flooding in Brassicaceae. Environmental and Experimental Botany 219, 105664.

Cook C.D.K. (1999) The number and kinds of embryo-bearing plants which have become aquatic: a survey. Perspectives in Plant Ecology, Evolution and Systematics 2, 79–102.

Dalle Carbonare L., Jiménez J.D.L.C., Lichtenauer S. & Van Veen H. (2023) Plant responses to limited aeration: Advances and future challenges. Plant Direct 7, e488.

Ellenberg, H. & Leuschner, C. (2010) Zeigerwerte der Pflanzen Mitteleuropas (Indicator values of vascular plants in Central Europe). In: Ellenberg, H. & Leuschner, C. (Eds.) Vegetation Mitteleuropas mit den Alpen (Vegetation of Central Europe including the Alps), 6th edition. Stuttgart: Ulmer.

Emms D.M. & Kelly S. (2019) OrthoFinder: phylogenetic orthology inference for comparative genomics. Genome Biology 20, 238.

Gan X., Hay A., Kwantes M., Haberer G., Hallab A., Ioio R.D.,… Tsiantis M. (2016) The *Cardamine hirsuta* genome offers insight into the evolution of morphological diversity. Nature Plants 2, 16167.

Gasch P., Fundinger M., Müller J.T., Lee T., Bailey-Serres J. & Mustroph A. (2016) Redundant ERF-VII Transcription Factors Bind to an Evolutionarily Conserved *cis*-Motif to Regulate Hypoxia-Responsive Gene Expression in Arabidopsis. The Plant Cell 28, 160– 180.

Grabherr M.G., Haas B.J., Yassour M., Levin J.Z., Thompson D.A., Amit I.,… Regev A. (2011) Full-length transcriptome assembly from RNA-Seq data without a reference genome. Nature Biotechnology 29, 644–652.

Haas B.J., Papanicolaou A., Yassour M., Grabherr M., Blood P.D., Bowden J.,… Regev A. (2013) De novo transcript sequence reconstruction from RNA-seq using the Trinity platform for reference generation and analysis. Nature Protocols 8, 1494–1512.

Hajheidari M., Wang Y., Bhatia N., Vuolo F., Franco-Zorrilla J.M., Karady M.,… Tsiantis M. (2019) Autoregulation of RCO by Low-Affinity Binding Modulates Cytokinin Action and Shapes Leaf Diversity. Current Biology 29, 4183–4192.e6.

Hartman S., Liu Z., Van Veen H., Vicente J., Reinen E., Martopawiro S.,… Voesenek L.A.C.J. (2019) Ethylene-mediated nitric oxide depletion pre-adapts plants to hypoxia stress. Nature Communications 10, 4020.

Hattori Y., Nagai K., Furukawa S., Song X.-J., Kawano R., Sakakibara H.,… Ashikari M. (2009) The ethylene response factors SNORKEL1 and SNORKEL2 allow rice to adapt to deep water. Nature 460, 1026–1030.

He D., Guo P., Gugger P.F., Guo Y., Liu X. & Chen J. (2018) Investigating the molecular basis for heterophylly in the aquatic plant *Potamogeton octandrus* (Potamogetonaceae) with comparative transcriptomics. PeerJ 6, e4448.

Howard H.W. & Lyon A.G. (1952) *Nasturtium officinale* R. Br. (*Rorippa nasturtium-aquaticum* (L.) Hayek). The Journal of Ecology 40, 228.

Hsu F.-C., Chou M.-Y., Chou S.-J., Li Y.-R., Peng H.-P. & Shih M.-C. (2013) Submergence Confers Immunity Mediated by the WRKY22 Transcription Factor in Arabidopsis. The Plant Cell 25, 2699–2713.

Ikematsu S., Umase T., Shiozaki M., Nakayama S., Noguchi F., Sakamoto T.,… Torii K.U. (2023) Rewiring of hormones and light response pathways underlies the inhibition of stomatal development in an amphibious plant *Rorippa aquatica* underwater. Current Biology 33, 543–556.e4.

Kantor A., Kučera J., Šlenker M., Breidy J., Dönmez A.A., Marhold K.,… Zozomová-Lihová J. (2023) Evolution of hygrophytic plant species in the Anatolia–Caucasus region: insights from phylogenomic analyses of *Cardamine* perennials. Annals of Botany 131, 585–600.

Kierzkowski D., Runions A., Vuolo F., Strauss S., Lymbouridou R., Routier-Kierzkowska A.-L.,… Tsiantis M. (2019) A Growth-Based Framework for Leaf Shape Development and Diversity. Cell 177, 1405–1418.e17.

Kim D., Paggi J.M., Park C., Bennett C. & Salzberg S.L. (2019) Graph-based genome alignment and genotyping with HISAT2 and HISAT-genotype. Nature Biotechnology 37, 907–915.

Kim J., Joo Y., Kyung J., Jeon M., Park J.Y., Lee H.G.,… Lee I. (2018) A molecular basis behind heterophylly in an amphibious plant, *Ranunculus trichophyllus*. PLOS Genetics 14, e1007208.

Koga H., Kojima M., Takebayashi Y., Sakakibara H. & Tsukaya H. (2021) Identification of the unique molecular framework of heterophylly in the amphibious plant *Callitriche palustris* L. The Plant Cell 33, 3272–3292.

Lee S.C., Mustroph A., Sasidharan R., Vashisht D., Pedersen O., Oosumi T.,… Bailey-Serres J. (2011) Molecular characterization of the submergence response of the *Arabidopsis thaliana* ecotype Columbia. New Phytologist 190, 457–471.

Leeggangers H.A.C.F., Rodriguez-Granados N.Y., Macias-Honti M.G. & Sasidharan R. (2023) A helping hand when drowning: The versatile role of ethylene in root flooding resilience. Environmental and Experimental Botany 213, 105422.

Li G., Yang J., Chen Y., Zhao X., Chen Y., Kimura S.,… Hou H. (2022) *SHOOT MERISTEMLESS* participates in the heterophylly of *Hygrophila difformis* (Acanthaceae). Plant Physiology 190, 1777–1791.

Liepman A.H. & Olsen L.J. (2003) Alanine Aminotransferase Homologs Catalyze the Glutamate:Glyoxylate Aminotransferase Reaction in Peroxisomes of Arabidopsis. Plant Physiology 131, 215–227.

Lin C., Zhang Z., Shen X., Liu D. & Pedersen O. (2024) Flooding-adaptive root and shoot traits in rice. Functional Plant Biology 51.

Ma X., Vanneste S., Chang J., Ambrosino L., Barry K., Bayer T.,… Van De Peer Y. (2024) Seagrass genomes reveal ancient polyploidy and adaptations to the marine environment. Nature Plants 10, 240–255.

Mommer L., Lenssen J.P.M., Huber H., Visser E.J.W. & De Kroon H. (2006) Ecophysiological determinants of plant performance under flooding: a comparative study of seven plant families. Journal of Ecology 94, 1117–1129.

Mommer L., Pons T.L., Wolters-Arts M., Venema J.H. & Visser E.J.W. (2005) Submergence-Induced Morphological, Anatomical, and Biochemical Responses in a Terrestrial Species Affect Gas Diffusion Resistance and Photosynthetic Performance. Plant Physiology 139, 497–508.

Müller J.T., Van Veen H., Bartylla M.M., Akman M., Pedersen O., Sun P.,… Mustroph A. (2021) Keeping the shoot above water – submergence triggers antithetical growth responses in stems and petioles of watercress (*Nasturtium officinale*). New Phytologist 229, 140–155.

Mustroph A., Lee S.C., Oosumi T., Zanetti M.E., Yang H., Ma K.,… Bailey-Serres J. (2010) Cross-Kingdom Comparison of Transcriptomic Adjustments to Low-Oxygen Stress Highlights Conserved and Plant-Specific Responses. Plant Physiology 152, 1484–1500.

Mustroph A., Zanetti M.E., Jang C.J.H., Holtan H.E., Repetti P.P., Galbraith D.W.,… Bailey-Serres J. (2009) Profiling translatomes of discrete cell populations resolves altered cellular priorities during hypoxia in Arabidopsis. Proceedings of the National Academy of Sciences 106, 18843–18848.

Nakamasu A., Nakayama H., Nakayama N., Suematsu N.J. & Kimura S. (2014) A Developmental Model for Branching Morphogenesis of Lake Cress Compound Leaf. PLoS ONE 9, e111615.

Nakayama H., Nakayama N., Seiki S., Kojima M., Sakakibara H., Sinha N. & Kimura S. (2014) Regulation of the KNOX-GA Gene Module Induces Heterophyllic Alteration in North American Lake Cress. The Plant Cell 26, 4733–4748.

Niessen M., Krause K., Horst I., Staebler N., Klaus S., Gaertner S.,… Peterhansel C. (2012) Two alanine aminotranferases link mitochondrial glycolate oxidation to the major photorespiratory pathway in Arabidopsis and rice. Journal of Experimental Botany 63, 2705–2716.

Nikolov L.A. & Tsiantis M. (2017) Using mustard genomes to explore the genetic basis of evolutionary change. Current Opinion in Plant Biology 36, 119–128.

Olsen J.L., Rouzé P., Verhelst B., Lin Y.-C., Bayer T., Collen J.,… Van De Peer Y. (2016) The genome of the seagrass *Zostera marina* reveals angiosperm adaptation to the sea. Nature 530, 331–335.

Pedersen O., Sauter M., Colmer T.D. & Nakazono M. (2021) Regulation of root adaptive anatomical and morphological traits during low soil oxygen. New Phytologist 229, 42– 49.

Rankenberg T., Van Veen H., Sedaghatmehr M., Liao C.-Y., Devaiah M.B., Stouten E.A.,… Sasidharan R. (2024) Differential leaf flooding resilience in *Arabidopsis thaliana* is controlled by ethylene signaling-activated and age-dependent phosphorylation of ORESARA1 activity. Plant Communications, 100848.

Reynoso M.A., Kajala K., Bajic M., West D.A., Pauluzzi G., Yao A.I.,… Bailey-Serres J. (2019) Evolutionary flexibility in flooding response circuitry in angiosperms. Science 365, 1291–1295.

Riber W., Müller J.T., Visser E.J.W., Sasidharan R., Voesenek L.A.C.J. & Mustroph A. (2015) The *Greening after Extended Darkness1* Is an N-End Rule Pathway Mutant with High Tolerance to Submergence and Starvation. Plant Physiology 167, 1616–1629.

Robinson M.D., McCarthy D.J. & Smyth G.K. (2010) edgeR: a Bioconductor package for differential expression analysis of digital gene expression data. Bioinformatics 26, 139–140.

Sasidharan R. & Voesenek L.A.C.J. (2015) Ethylene-Mediated Acclimations to Flooding Stress. Plant Physiology 169, 3–12.

Sasidharan R., Hartman S., Liu Z., Martopawiro S., Sajeev N., Van Veen H.,… Voesenek L.A.C.J. (2018) Signal Dynamics and Interactions during Flooding Stress. Plant Physiology 176, 1106–1117.

Sasidharan R., Mustroph A., Boonman A., Akman M., Ammerlaan A.M.H., Breit T.,… Van Tienderen P.H. (2013) Root Transcript Profiling of Two *Rorippa* Species Reveals Gene Clusters Associated with Extreme Submergence Tolerance. Plant Physiology 163, 1277–1292.

Shimizu-Inatsugi R., Terada A., Hirose K., Kudoh H., Sese J. & Shimizu K.K. (2017) Plant adaptive radiation mediated by polyploid plasticity in transcriptomes. Molecular Ecology 26, 193–207.

Smit M.E. & Bergmann D.C. (2023) The stomatal fates: Understanding initiation and enforcement of stomatal cell fate transitions. Current Opinion in Plant Biology 76, 102449.

Stift M., Luttikhuizen P.C., Visser E.J.W. & Van Tienderen P.H. (2008) Different flooding responses in *Rorippa amphibia* and *Rorippa sylvestris*, and their modes of expression in F _1_ hybrids. New Phytologist 180, 229–239.

Sucher J., Mbengue M., Dresen A., Barascud M., Didelon M., Barbacci A. & Raffaele S. (2020) Phylotranscriptomics of the Pentapetalae Reveals Frequent Regulatory Variation in Plant Local Responses to the Fungal Pathogen *Sclerotinia sclerotiorum*. The Plant Cell 32, 1820–1844.

Summers J.E., Voesenek L.A.C.J., Blom C.W.P.M., Lewis M.J. & Jackson M.B. (1996) *Potamogeton pectinatus* Is Constitutively Incapable of Synthesizing Ethylene and Lacks 1-Aminocyclopropane-1-Carboxylic Acid Oxidase. Plant Physiology 111, 901– 908.

Usadel B., Bläsing O.E., Gibon Y., Retzlaff K., Höhne M., Günther M. & Stitt M. (2008) Global Transcript Levels Respond to Small Changes of the Carbon Status during Progressive Exhaustion of Carbohydrates in Arabidopsis Rosettes. Plant Physiology 146, 1834– 1861.

Van Veen H. & Sasidharan R. (2021) Shape shifting by amphibious plants in dynamic hydrological niches. New Phytologist 229, 79–84.

Van Veen H., Akman M., Jamar D.C.L., Vreugdenhil D., Kooiker M., Van Tienderen P.,… Sasidharan R. (2014) Group VII E thylene R esponse F actor diversification and regulation in four species from flood-prone environments. Plant, Cell & Environment 37, 2421–2432.

Van Veen H., Mustroph A., Barding G.A., Vergeer-van Eijk M., Welschen-Evertman R.A.M., Pedersen O.,… Sasidharan R. (2013) Two *Rumex* Species from Contrasting Hydrological Niches Regulate Flooding Tolerance through Distinct Mechanisms. The Plant Cell 25, 4691–4707.

Van Veen H., Vashisht D., Akman M., Girke T., Mustroph A., Reinen E.,… Sasidharan R. (2016) Transcriptomes of eight *Arabidopsis thaliana* accessions reveal core conserved, genotype- and organ-specific responses to flooding stress. Plant Physiology, pp.00472.2016.

Vashisht D., Hesselink A., Pierik R., Ammerlaan J.M.H., Bailey-Serres J., Visser E.J.W.,… Sasidharan R. (2011) Natural variation of submergence tolerance among *Arabidopsis thaliana* accessions. New Phytologist 190, 299–310.

Voesenek L.A.C.J. & Bailey-Serres J. (2013) Flooding tolerance: O2 sensing and survival strategies. Current Opinion in Plant Biology 16, 647–653.

Voesenek L.A.C.J. & Sasidharan R. (2013) Ethylene – and oxygen signalling – drive plant survival during flooding. Plant Biology 15, 426–435.

Voesenek L.A.C.J., Pierik R. & Sasidharan R. (2015) Plant Life without Ethylene. Trends in Plant Science 20, 783–786.

Weits D.A., Giuntoli B., Kosmacz M., Parlanti S., Hubberten H.-M., Riegler H.,… Licausi F. (2014) Plant cysteine oxidases control the oxygen-dependent branch of the N-end-rule pathway. Nature Communications 5, 3425.

Wittig P.R., Ambros S., Müller J.T., Bammer B., Álvarez-Cansino L., Konnerup D.,… Mustroph A. (2021) Two *Brassica napus* cultivars differ in gene expression, but not in their response to submergence. Physiologia Plantarum 171, 400–415.

Wu J., Zhao H.-B., Yu D. & Xu X. (2017) Transcriptome profiling of the floating-leaved aquatic plant *Nymphoides peltata* in response to flooding stress. BMC Genomics 18, 119.

Xu K., Xu X., Fukao T., Canlas P., Maghirang-Rodriguez R., Heuer S.,… Mackill D.J. (2006) Sub1A is an ethylene-response-factor-like gene that confers submergence tolerance to rice. Nature 442, 705–708.

Yeung E., Van Veen H., Vashisht D., Sobral Paiva A.L., Hummel M., Rankenberg T.,… Sasidharan R. (2018) A stress recovery signaling network for enhanced flooding tolerance in *Arabidopsis thaliana*. Proceedings of the National Academy of Sciences 115.

Young M.D., Wakefield M.J., Smyth G.K. & Oshlack A. (2010) Gene ontology analysis for RNA-seq: accounting for selection bias. Genome Biology 11, R14.

