## Supplemental Figure for "Phylotranscriptomics provides a treasure trove of flood tolerance mechanisms in the Cardamineae tribe"

### SUPPLEMENTAL METHODS

#### Transcriptome assembly and orthology analysis

The *de novo* assembly of the transcriptomes for *C. pratensis*, *R. palustris* and *R. sylvestris* were done with Trinity version 2.6.6 (Grabherr et al. 2011, Haas et al. 2013), with the default Kmer size of 25, a minimal Kmer coverage of 2 and with the stranded option. For *C. hirsuta* a transcriptome was assembled guided by the genome from Gan et al. (2016). To this end first a spliced alignment to the genome was done by HISAT2 with a maximum intron length of 6000 and increased anchor requirements by the "dta" option to detect novel splice sites (Kim et al. 2019). Additionally, only concordant hits meeting the strandedness requirements were considered. The resulting alignments were used by Trinity v2.6.6 with the default settings to produce a de novo transcriptome.

Trinity outputs multiple isoforms from an assembly graph. Only one representative transcript isoform was used to detect orthogroups. This primary transcript was selected based on quality ranking where long transcripts with many mapped reads were preferred and where, based on a discontiguous blast search to Araport 11 cDNA, there was the lowest evidence of a chimeric assembly. The primary transcripts of all assembled transcriptomes, primary isoforms from *A. thaliana* (Araport11) and from *R. islandica* (Phytozome) were subjected to an all-vs-all discontiguous megablast (Camacho et al. 2009). Next the blast output format was adjusted to provide compatibility with Orthofinder (Emms and Kelly 2019). Orthofinder was run with various inflation parameters to find optimal orthogroup separation (Supplemental Figure S5). An inflation score of 1.6 was used.

#### Differential expression and Gene Ontology enrichment

To calculate the fold changes and differential expression, only genes or orthogroups with more than 10 reads in at least 3 samples were considered. Orthogroup read count was the sum of all the counts of the genes in the orthogroup. EdgeR was used to calculate the fold changes and significance (Robinson et al. 2010). The submergence response after 24 h, 48 h and the difference between the time points (time*treatment interaction) were estimated with a separate full factorial model for each species. The main effect of flooding was estimated similarly, but using an additive model.

Between species effects were identified similarly but with an additional phylogeny or tolerance effect. We repeatedly applied a model with either a factor for species (5 levels: Atha, Chir, Cpra, Rpal and Rsyl), a factor for genus (3 levels: Arabidopsis, Cardamine and Rorippa), or a factor for tolerance and tribe (2 levels: *A. thaliana* and Cardamineae). Additionally, we used a factor for the tolerance grouping with 3 levels, namely sensitive (Atha), moderately tolerant (Chir) and very tolerant (Cpra, Rpal, Rsyl). In the case of genus, tribe and the tolerance groups, the design was nested such that variation between and within species and the factor levels could be partitioned accordingly. The mean species effect was estimated with the species factor, but then in an additive design rather than a factorial design.

Deviation from the mean response was defined as the log2FC from a species minus the mean log2FC.

The orthogroup selection criteria (Pvalue and Fold Change) for the varied analyses are provided in the corresponding figure and results. Orthogroup clustering was performed in R (v4.3.1) by distance calculation with the dist() function with Euclidean method and subsequent clustering with hclust() using the “ward.D2” method. Gene Ontology enrichment was performed using the GOseq package (Young et al. 2010) using manually entered GOterm annotations. Orthogroup GOterms were annotated based on the *A. thaliana* genes present. In case of multiple *A. thaliana* genes, the GO terms would be combined. In the absence of an *A. thaliana* gene, annotation was by best blast hit to *A. thaliana* (Eval < 1e-40). *A. thaliana* annotation was obtained from the 'org.At.tair.db' (Carlson 2019a) and 'GO.db' libraries (Carlson 2019b).

**Summary of the used factors and their levels**

Time: 24 hours and 48 hours
Treatment: control and submergence
Species: *A. thaliana* (Atha), *C. hirsuta* (Chir), *C. pratensis* (Cpra), *R. palustris* (Rpal), and *R. sylvestris* (Rsyl)
Genera: Arabidopsis (Ara), Cardamine (Car), Rorippa (Ror)
Tribe: Cammelineae (*A. thaliana*), and Cardamineae (*C. hirsuta*, *C. pratensis*, *R. palustris*, *R. sylvestris*)
Tolerance groups: sen/sensitive (*A. thaliana*), mod/moderately tolerant (*C. hirsuta*), tol/tolerant (*C. pratensis*, *R. palustris*, *R. sylvestris*)

**Summary and abbreviations of tested effects**

*Experimental effects* (Supplemental data S2, S3)
sub1: submergence effect at 24 hours
sub2: submergence effect at 48 hours
TxT: time*treatment effect, i.e. sub2 minus sub1
subM: main submergence effect, i.e. the mean response over both timepoint

*Experimental effects* (Supplemental data S5)
A: average submergence effect
sub1: submergence effect at 24 hours
sub2: submergence effect at 48 hours
subM: main submergence effect, i.e. the mean response over both timepoint

*Difference in response between species* (Supplemental data S5)
SxTrsub1, SxTrsub2, SxTrsubM:
species*treatment interaction effect
Tests whether species vary in their response to submergence for 24h/48h/Main

*GxTrsub1, GxTrsub2, GxTrsubM* (Supplemental data S5)
genus*treatment interaction effect
Tests whether genera vary in their response to submergence for 24h/48h/Main
The model fits identical responses within a genus
Genus and species are nested accordingly

*TolxTr1, TolxTr2, TolxTrsubM* (Supplemental data S5)
tribe*treatment interaction effect, for 24h/48h/Main
i.e. the response of the Cardamineae minus the response of the Cammelineae
The model fits identical responses within a tribe
Species and tribe are nested accordingly

*TGxTrsub1, TGxTrsub2, TGxTrsubM:*
tolerance-group*treatment effect, for 24h/48h/Main

*GO enrichment definitions:*
numDEInCat: number of OGs belonging to a GO-category that are differentially expressed or in a cluster of interest
notDEInCat: number of OGs belonging to a GO-category that are NOT differentially expressed or NOT in a cluster of interest
numDEOutCat: number of OGs noy belonging to a GO-category that are differentially expressed or in a cluster of interest
notDEOutCat: number of OGs not belonging to a GO-category that are NOT differentially expressed or NOT in a cluster of interest
Oddsratio: quantifies the magnitude of enrichment by taking the ratio of the odds to belong to a GO-category when differentially expressed or in a cluster, and belong to a GO-category when NOT differentially expressed or NOT in a cluster i.e. (numDEInCat/numDEoutCat) / (notDEInCat/notDEoutCat)
over_represented_pvalue: significance of enrichment

**SUPPLEMENTAL FIGURES**

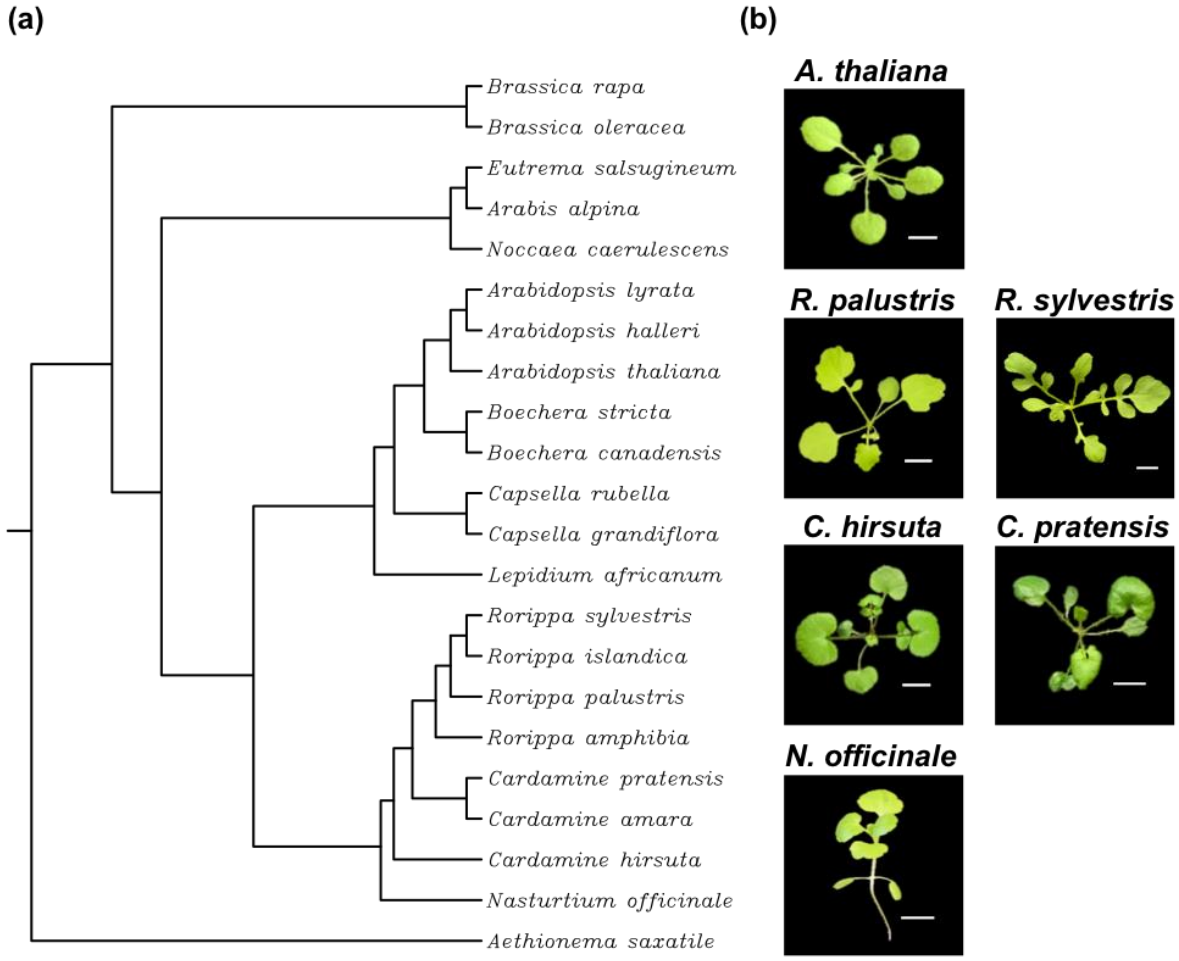


**Figure S1.** Phylogenetic and phenotypic characterization of selected Brassicaceae. (a) Phylogenetic tree based on ITS1 sequences (*internal transcribed spacer 1*), obtained from NCBI. The tree was constructed following a multiple sequence alignment with ClustalW. Representative images of the studied species. (b) Representative images of the plants (3 – 4 weeks old) used in this study and in Müller et al. 2021. Scale bar = 1 cm.

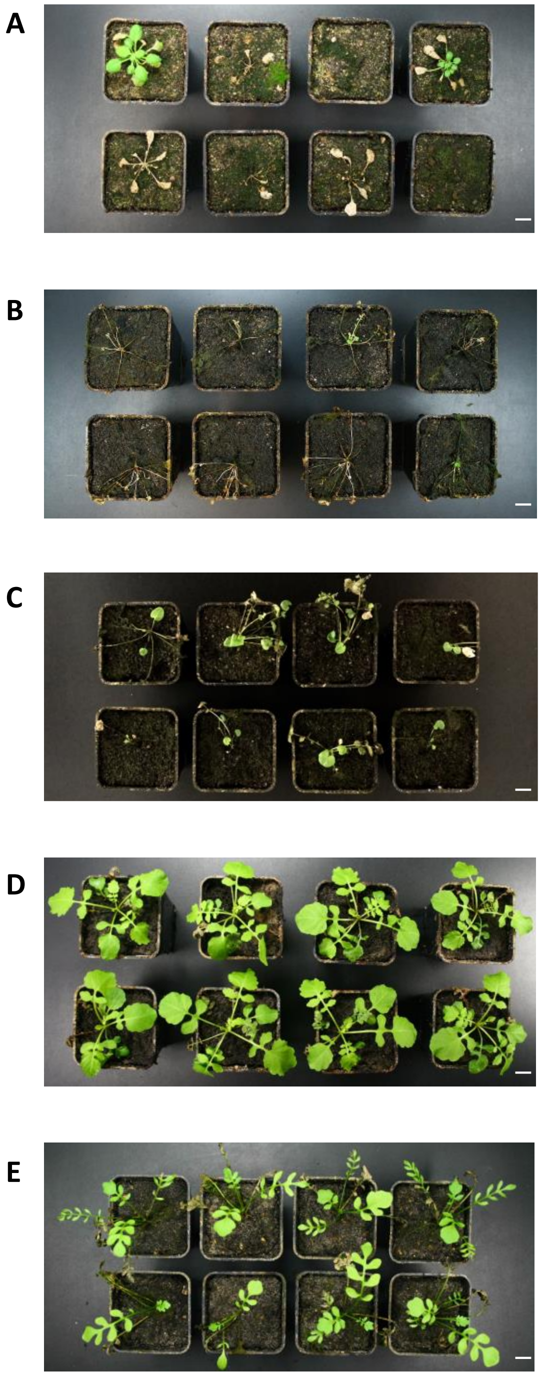


**Figure S2.** Pictures of plants after submergence stress and a two-week-recovery phase. Quantification of data is shown in Figure 1A. (a) *A. thaliana*, 3.5 weeks of submergence; (b) *C. hirsuta*, 8 weeks of submergence; (c) *C. pratensis*, 10 weeks of submergence; (d) *R. palustris*, 10 weeks of submergence; (e) *R. sylvestris*, 10 weeks of submergence.

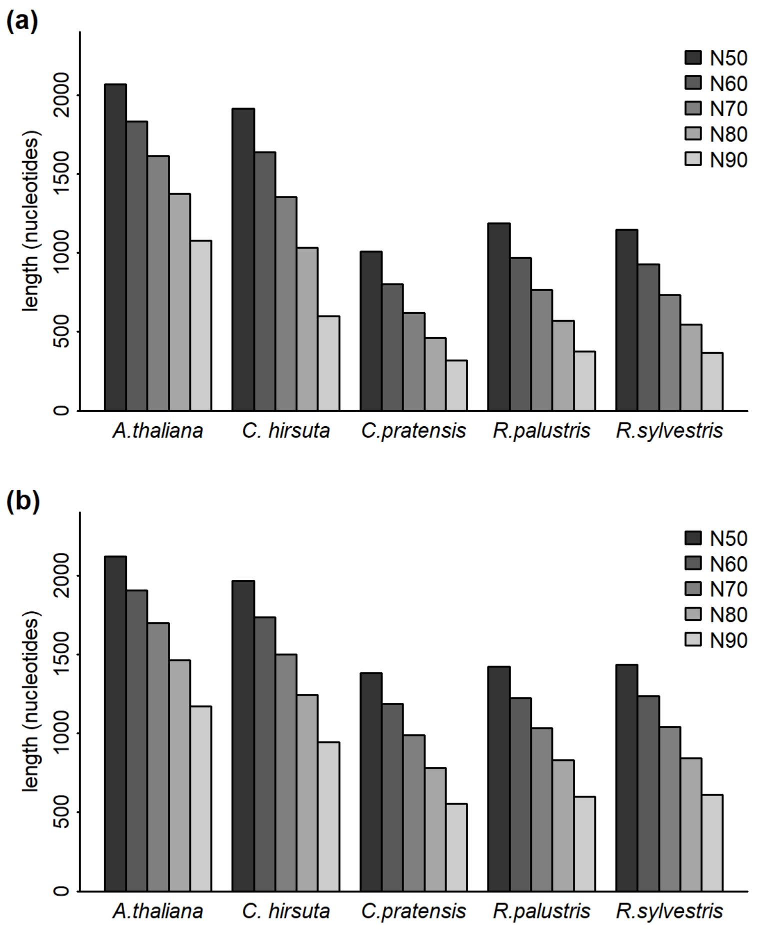


**Figure S3.** N50 to N90 statistics of all assembled transcripts and the *A. thaliana* transcriptome (a) and the assembled transcripts that represent the universal orthogroups defined in Figure 2 (b). N50 is the length of the shortest transcript where all longer and equal length transcripts cover 50% of the assembled nucleotides.

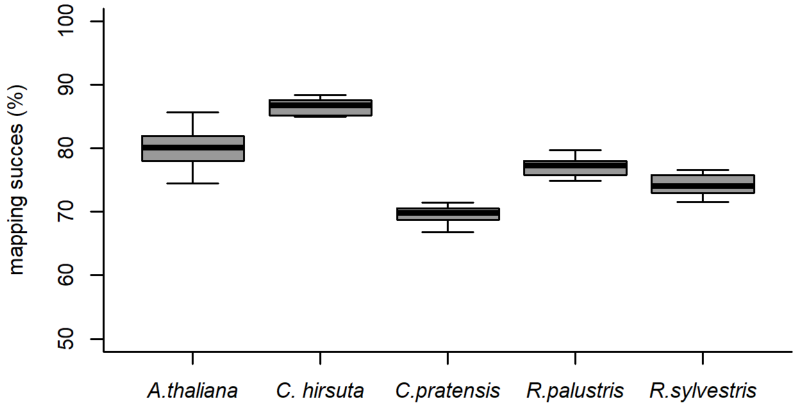


**Figure S4.** Percentage of reads that successfully mapped to the reference transcriptome.

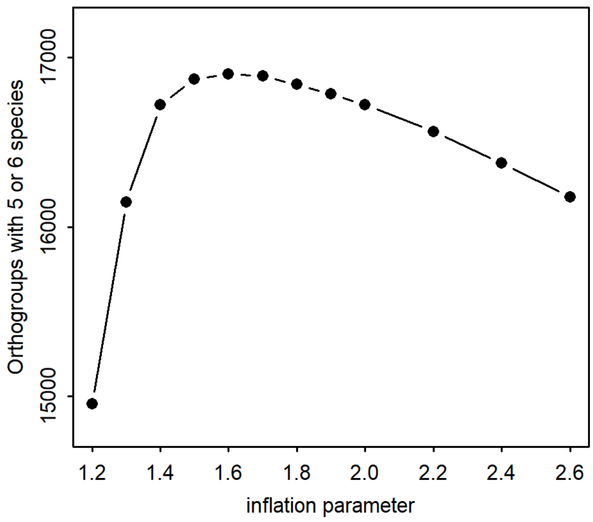


**Figure S5.** To identify groups of orthologs with Orthofinder (Emms and Kelly 2019) a graph network is created based on an all-vs-all blast. Subsequently the MCL clustering algorithm separates the graph network into smaller highly connected graphs, which results in distinct orthogroups. The inflation parameter determines to what extent graphs are split (high inflation value) or lumped together (low inflation value). To obtain the smallest possible orthogroup, that still was represented by all species we ran Orthofinder with a range of inflation parameters to obtain the optimal result.

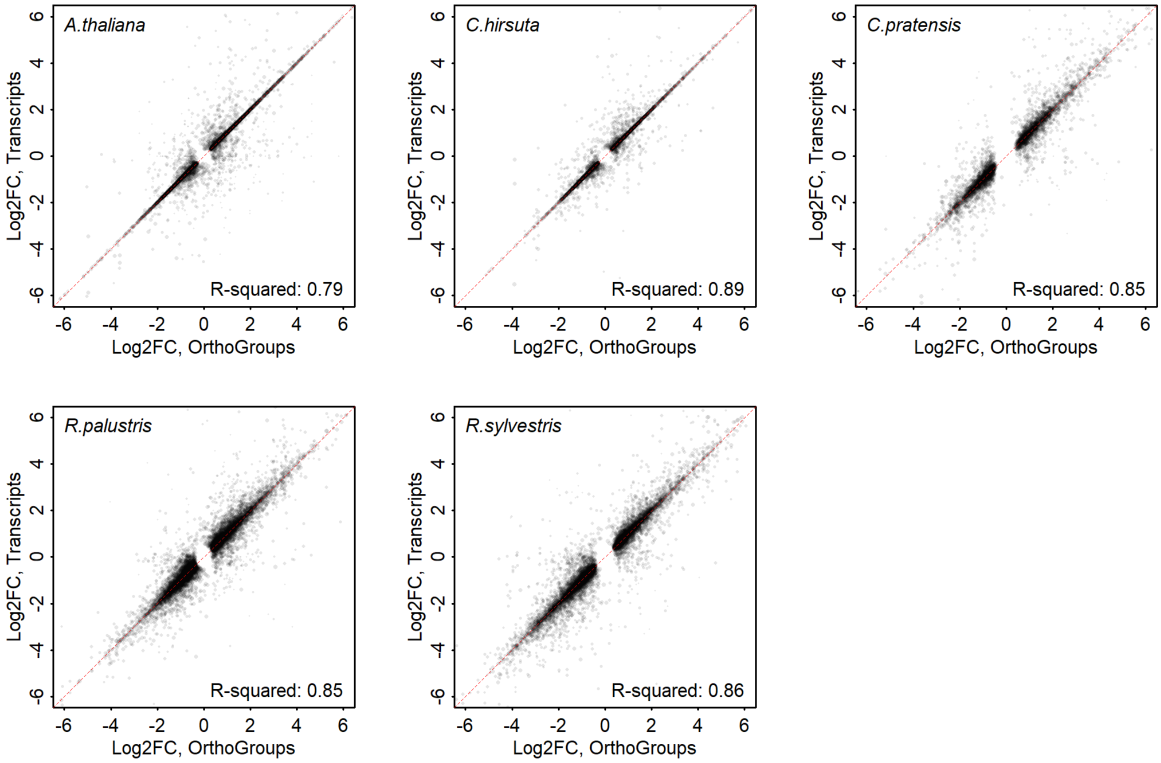


**Figure S6.** The extent to which the fold change of a transcript can be predicted by the fold change of the orthogroup. The dashed red line depicts y = x, which is also the model on which the $R^2$ is based. Here $R^2 = 1 - (SS_{residuals}/SS_{total})$. All transcripts from the universal orthogroups that were differentially expressed at either 24 or 48 h are shown, therefore some transcripts are present twice, for their 24 h and their 48 h estimate.

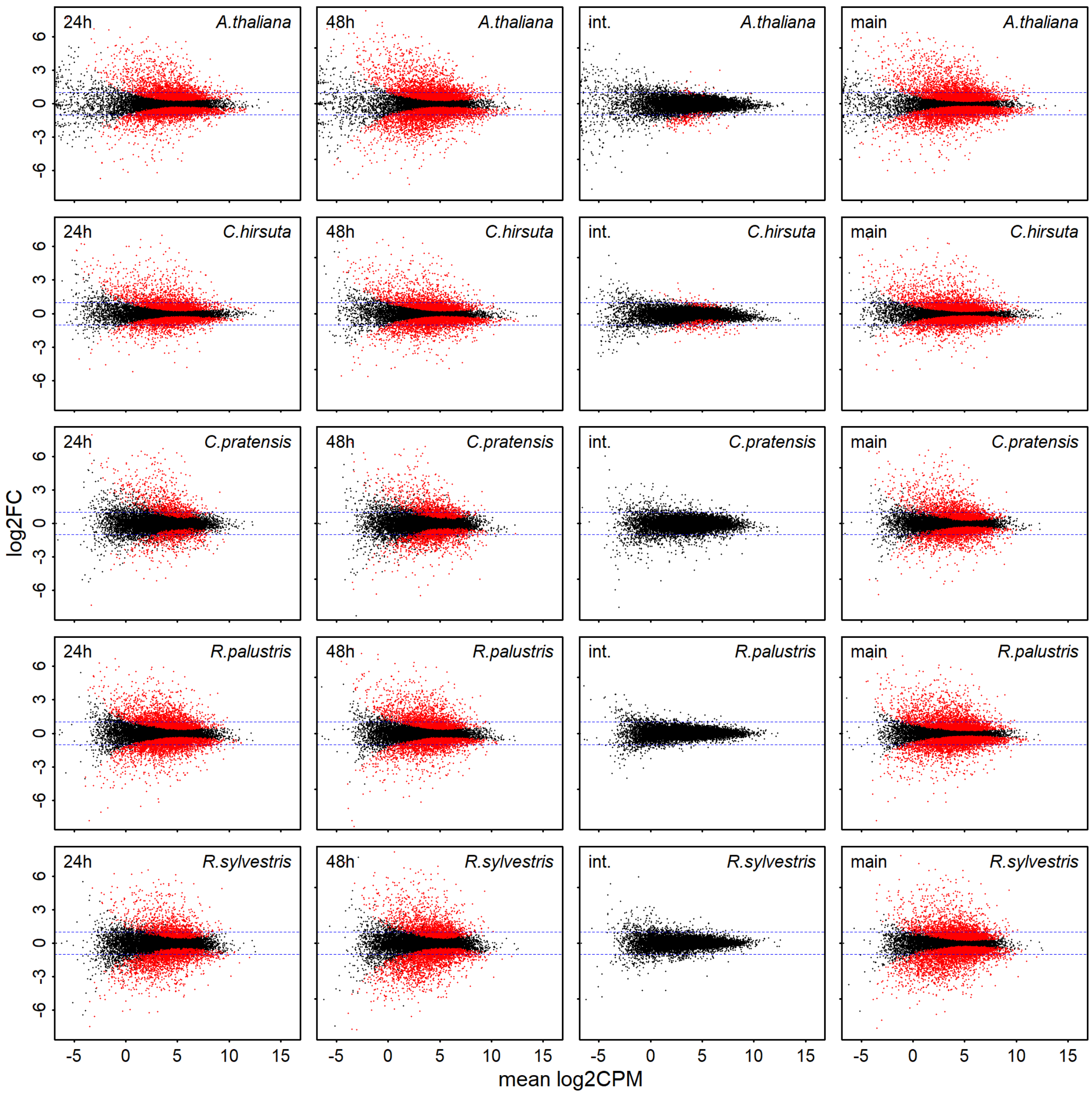


**Figure S7.** The fold change of orthogroups compared to their mean sequencing depth across all samples (CPM, counts per million). The orthogroups shown in red were significantly regulated ($P_{adj.} < 0.01$). Here int. considers the time*treatment interaction effect and main the main effect of flooding in an additive model with time and treatment as factors. The dashed blue line indicates |log2FC| = 1.

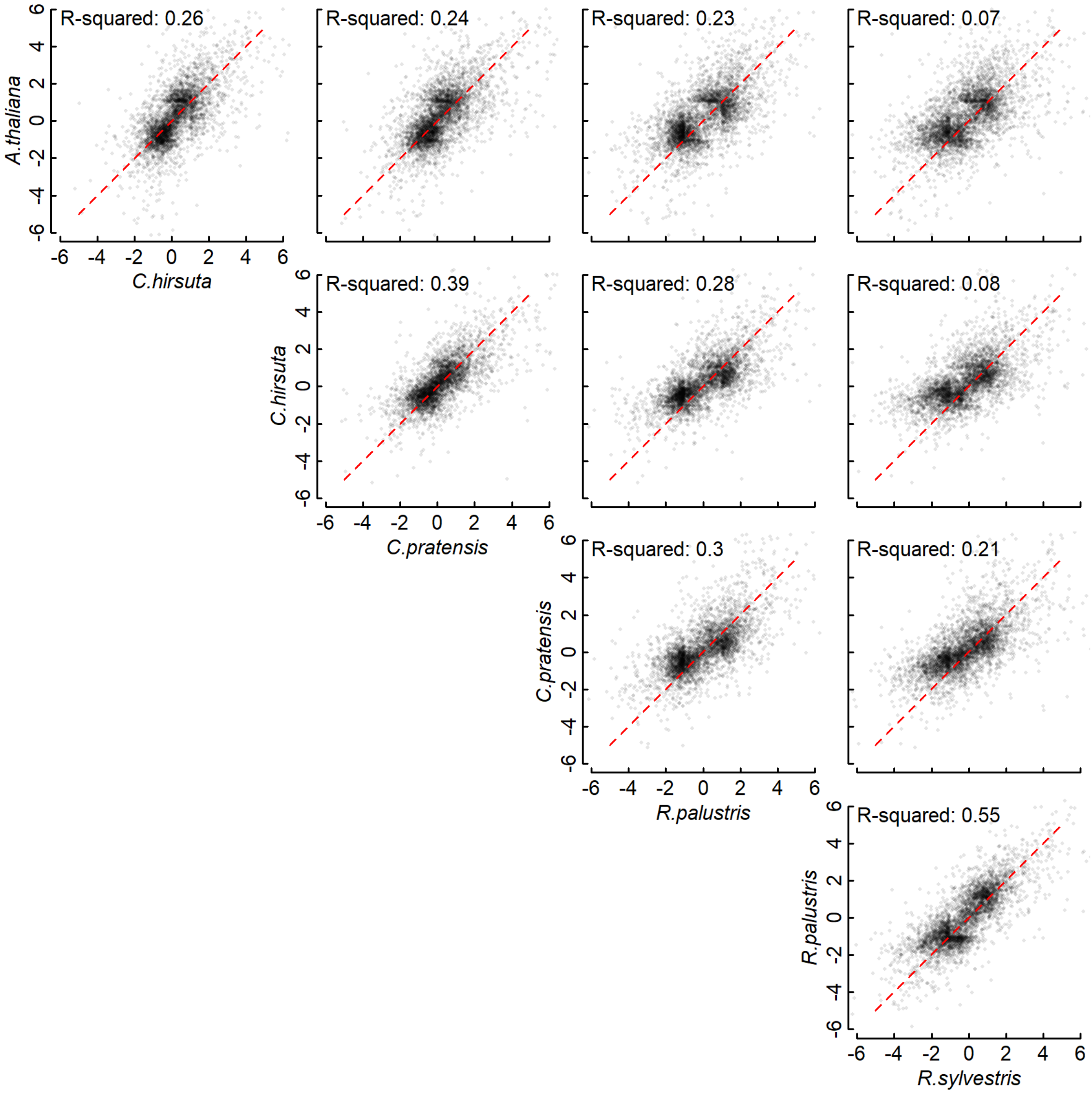


**Figure S8.** Pairwise comparison of 24 h flood responses between all species combinations. Orthogroups that were differentially expressed ($P_{adj.} < 0.001$, $|log_2FC| > 1$) in at least one species at the 24 h timepoint are shown in all graphs. The dashed red line depicts y = x, which is also the model on which the $R^2$ is based. Here $R^2 = 1 - (SS_{residuals\text{-}speciesA} + SS_{residuals\text{-}speciesB}) / (SS_{total\text{-}speciesA} + SS_{total\text{-}speciesA})$.

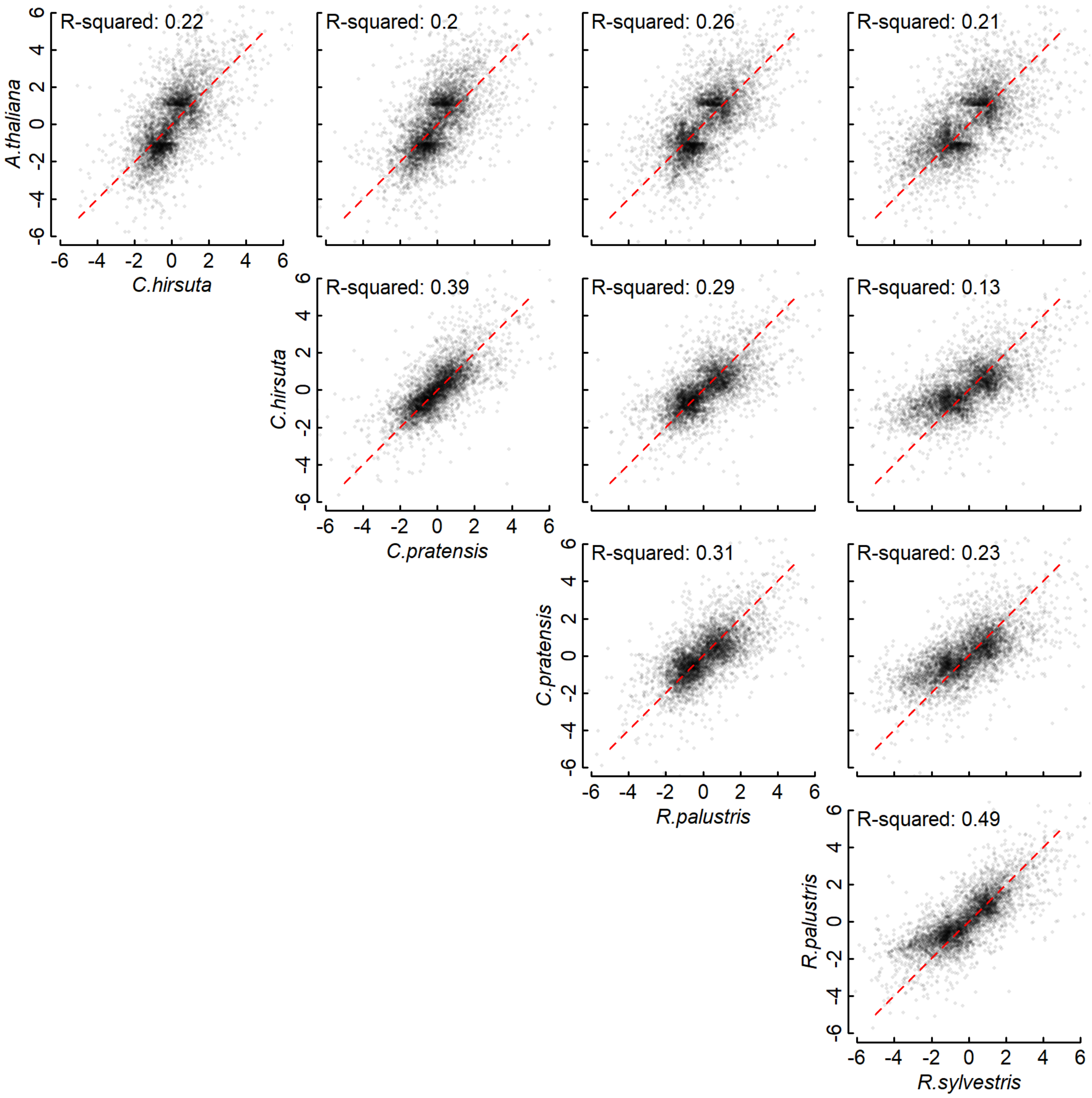


**Figure S9.** Pairwise comparison of 48 h flood responses between all species combinations. Orthogroups that were differentially expressed ($P_{adj.} < 0.001$, $|log_2FC| > 1$) in at least one species at the 48 h timepoint are shown in all graphs. The dashed red line depicts y = x, which is also the model on which the $R^2$ is based. Here $R^2 = 1 - (SS_{residuals\text{-}speciesA} + SS_{residuals\text{-}speciesB}) / (SS_{total\text{-}speciesA} + SS_{total\text{-}speciesA})$.
