## Supplementary material for "Phylotranscriptomics provides a treasure trove of flood tolerance mechanisms in the Cardamineae tribe": Abbreviations.txt

1. Supplemental datasets
2. used factors and their levels
3. tested effects
4. GO enrichment data


####
1. Supplemental datasets

S1:  Assignment of genes to orthogroups (OGs)
S2a: Fold changes of genes, members of universal (OGs)
S2b: Fold changes of genes, not a member of universal (OGs)
S3a: Fold changes of universal (OGs)
S3b: Fold changes of species specific Orthogroups
S4a: Gene Ontology enrichment among downregulated OGs
S4b: Gene Ontology enrichment among upregulated OGs
S5:  Between Species effects
S6:  Clustermembership of OGs
S7a: Gene Ontology enrichment of clusters without species effect
S7b: Gene Ontology enrichment of clusters with genus effect
S7c: Gene Ontology enrichment of clusters associated with tolerance
S8:  OGs used in targeted analysis


####
2. used factors and their levels

time
  24 hours
  48 hours

treatment
  control
  submergence

Species
  Atha: A. thaliana
  Chir: C. hirsuta
  Cpra: C. pratensis
  Rpal: R. palustris
  Rsyl: R. sylvestris

Genera:
  Ara: Arabidopsis
  Car: Cardamine
  Ror: Rorippa

Tribe:
  Cammelineae: A. thaliana
  Cardamineae: C.hirsuta, C.pratensis,R.palustris,R.sylvestris

Tolerance groups
  sen: sensitive; A.thaliana
  mod: moderately tolerant; C.hirsuta
  tol: tolerant; C.pratensis, R.palustris, R.sylvestris


####
3. tested effects

Experimental effects
  sub1: submergence effect at 24 hours
  sub2: submergence effect at 48 hours
  TxT: time*treatment effect, i.e. sub2 minus sub1
  subM: main submergence effect, i.e. the mean response over both timepoint

Difference in response between species

  SxTrsub1, SxTrsub2, SxTrsubM: 
    species*treatment interaction effect
    Tests whether species vary in their response to submergence for 24h/48h/Main

  GxTrsub1, GxTrsub2, GxTrsubM:
    genus*treatment interaction effect
    Tests whether genera vary in their response to submergence for 24h/48h/Main.
    The model fits identical responses within a genus
    genus and species are nested accordingly

  TolxTr1, TolxTr2, TolxTrsubM:
    tribe*treatment interaction effect, for 24h/48h/Main
    i.e. the response of the Cardamineae minus the response of the Cammelineae
    The model fits identical responses within a tribe
    Species and tribe are nested accordingly

  TGxTrsub1, TGxTrsub2, TGxTrsubM:
    tolerance-group*treatment effect, for 24h/48h/Main

####
4. GO enrichment data

numDEInCat:
  number of OGs belonging to a GO-category that are
  differentially expressed or in a cluster of interest
notDEInCat:
  number of OGs belonging to a GO-category that are
  NOT differentially expressed or NOT in a cluster of interest
numDEOutCat:
  number of OGs noy belonging to a GO-category that are
  differentially expressed or in a cluster of interest
notDEOutCat:
  number of OGs not belonging to a GO-category that are
  NOT differentially expressed or NOT in a cluster of interest

Oddsratio:
  quantifies the magnitude of enrichment by taking the ratio of the odds to 
     belong to a GO-category when differentially expressed or in a cluster, 
     and belong to a GO-category when NOT differentially expressed or NOT in a cluster
  i.e. (numDEInCat/numDEoutCat) / (notDEInCat/notDEoutCat)

over_represented_pvalue:
  significance of enrichment
